# Costless metabolic secretions as drivers of interspecies interactions in microbial ecosystems

**DOI:** 10.1101/300046

**Authors:** Alan R. Pacheco, Mauricio Moel, Daniel Segrè

## Abstract

Metabolic exchange can mediate beneficial interactions among microbes, helping explain diversity in microbial communities. These interactions are often assumed to involve a fitness cost, prompting questions on how cooperative phenotypes can be stable and withstand the emergence of cheaters. Here we use genome-scale models of metabolism to investigate whether a radically different scenario, the pervasive release of “costless” metabolites (i.e. those that cause no fitness cost to the producing organism), can serve as a prominent mechanism for inter-microbial interactions. By carrying out over 1 million pairwise growth simulations for 14 microbial species in a combinatorial assortment of environmental conditions, we find that there is indeed a large space of metabolites that can be secreted at no cost, which can generate ample cross-feeding opportunities. In addition to providing an atlas of putative costless interdependencies, our modeling also demonstrates that oxygen availability significantly enhances mutualistic interactions by providing more opportunities for metabolic exchange through costless metabolites, resulting in an over-representation of specific ecological network motifs. In addition to helping explain natural diversity, we show how the exchange of costless metabolites can facilitate the engineering of stable synthetic microbial consortia.

## INTRODUCTION

The astonishing number of microbial species observed in nature ^1–3^ seems to contradict classical ecological theory, which predicts far less biodiversity in many nutrient-poor environments ^4,5^. A variety of different explanations have been proposed as possible solutions to this inconsistency, including resource partitioning ^6^, differential nutrient use ^7^, spatial segregation ^8^, and metabolic cross-feeding ^9–11^. In environments poor in resources, cross-feeding has been shown to enhance the capacity of microbes to survive, either through the secretion of valuable compounds ^12–14^, or by maintaining thermodynamic gradients necessary for continued metabolism ^15^. Despite their prevalence, it is not clear how these cooperative phenotypes emerge, as they often involve the exchange of metabolites that are costly for the producer. This apparent altruism introduces the potential for the rise of cheating organisms that do not contribute common goods but still benefit metabolically from others, challenging community stability ^16^. Previous studies have addressed this dilemma in different ways, either by suggesting fundamental boundaries to the feasibility of costly cross-feeding based on theoretical considerations ^17^, or by establishing balances between production costs and reciprocation benefits in specific communities and environments ^18–20^. However, it is not fully understood whether costly exchanges can account for the degree of biodiversity observed in nature, as the conditions necessary for the rise and maintenance of these costly interdependencies may not frequently manifest themselves.

We therefore ask whether a radically different interaction mechanism, one that hinges on organisms secreting metabolic products at no cost to their own fitness, may be prevalent enough in the microbial world to help explain the abundance of cross-feeding opportunities. Key to this mechanism, which we will term “costless,” is the emergence of community benefits as a product of otherwise selfish acts by individual microbial species. This phenomenon has been explored in a macroecological context ^21–23^ and can be illustrated by the example of a vulture consuming the remains of a lion kill. Here, the lion gains nutritional benefit from its hunt and leaves behind scraps of food that are in turn eaten by the vulture. In this way, though the lion did not expend energy to facilitate access to food explicitly for the vulture, it did unintentionally contribute to the vulture’s success through its own selfish action ^24^. It is known that, in the microbial world, metabolic waste products secreted at no cost to the producing organism (e.g. *E. coli* secreting acetate under limited oxygen) can serve to support other species ^13^. However, it is not obvious whether such behavior extends beyond a few fermentation byproducts. Moreover, little information exists on how costless secretions vary across microbial species and growth media composition. Most importantly, even if the metabolites secreted by an organism under a given condition were to be known, it still would be difficult to ascertain whether such byproducts would be likely to enable or enhance growth of other species.

In this study, we use computational metabolic modeling to quantify the magnitude of environmental modification brought about by costless metabolite secretion, as well as the interspecies interactions that can arise from this type of exchange. In a microbial analog to the lion-vulture interaction, we seek to understand how metabolites released as a product of selfish action by individual species yield unintended benefits to partner organisms, resulting in emergent interspecies cooperation. Based on this framework, we present a computational pipeline based on flux balance analysis (FBA) ^25^ that predicts the growth phenotypes and cooperative interactions mediated by costless metabolites for 14 microbial species under a large combinatorial set of environmental conditions. In this way, we obtain a global view of cross-feeding opportunities that can mediate the emergence of cooperation and the maintenance of biodiversity in natural communities. In addition, we complement our metabolic modeling with a dynamical modeling framework to understand whether costless secretions on their own can promote long-term stability in model synthetic microbial communities. While the present work focuses entirely on putative secretions and interactions predicted computationally, we wish to highlight that we restricted our analysis to microbes associated with high quality, manually curated (and therefore in most cases individually tested) *in silico* models and that in many cases, specific predictions can be shown to be consistent with previously established empirical knowledge. For the most part, however, the current analysis should be viewed as the exploration of a large space of stoichiometrically possible costless interactions (inscrutable to such an extent at the experimental level), whose global patterns could motivate and inform future experimental and theoretical endeavors.

## RESULTS

### Metabolite secretions can be costly or costless, depending on environmental context

Understanding whether or not the secretion of a specific metabolite by a given organism is associated with a decrease in fitness (interpreted here as growth rate) is difficult to assess experimentally, but can be readily addressed using genome-scale models of metabolism (see Methods). For example, it is possible to impose the secretion of a given compound at a given rate, and then ask whether this constraint is expected to cause a reduction in growth. A small set of simulations of this kind for a single organism (Fig. S1) exemplifies the broad spectrum of possible outcomes: based on the specific carbon sources, different metabolites can be produced, sometimes at the expense of growth capacity, other times with no apparent effect (neutral), or even to its benefit. Due to the basic assumptions of the genome-scale models we employed (especially the maximization of growth as the objective function) we know that these last two kinds of secretions are compatible (or even necessary) for metabolism to operate at maximal efficiency. We will refer to these beneficial or neutral secretions as ‘costless.’

### Secretion of costless metabolites leads to substantive environmental enrichment

Having illustrated in an individual case how metabolite secretion costs can strongly depend on carbon sources, we sought to map the prevalence of costless secretions across a broad set of organisms and environments, as well as the chance that such secreted metabolites could mediate cross-feeding. We carried out a total of 1,051,596 unique *in silico* simulations, each with two organisms *i* and *j* from a set of 14 genome-scale metabolic models and two carbon sources *α* and *β* from a set of 108 compounds (Figure 1, Supplementary Information 1, 2). Each simulation is conducted as an iterative process that simulates a coculture experiment, and is uniquely defined by the organisms involved, the carbon sources provided, and the availability of oxygen. At each iteration, we used FBA to determine the ability of each organism to grow on the current medium (see Methods, and Fig. 1a). As an outcome of this calculation, we also obtained information about the set of metabolites predicted to be spontaneously (i.e. costlessly) secreted by each microbe. If at the first iteration (*c* = 1) at least one *in silico* organism was able to grow on the carbon sources provided, all costless metabolites were added to the medium for the next iteration. This process was repeated until no new metabolites were produced. The final iteration *c* before attaining this steady state is defined as *c_s_*. Upon running these iteration simulations for all possible combinations of species and environments and selecting only the cases in which both species grew, we obtained distributions for the value of *c_s_* (Figure 2a). A large number of cases reached a steady state after only one iteration (92% of cases with oxygen, and in 82% without oxygen). This could in principle be due to a complete lack of costless secretions at the first iteration. However, as demonstrated by Fig. 2b, the skewness in the distribution of *c_s_* is better explained by the alternative hypothesis that organisms do secrete multiple byproducts in the first iteration, but these byproducts contribute weakly to additional secretions in subsequent iterations.

**Figure 1.**
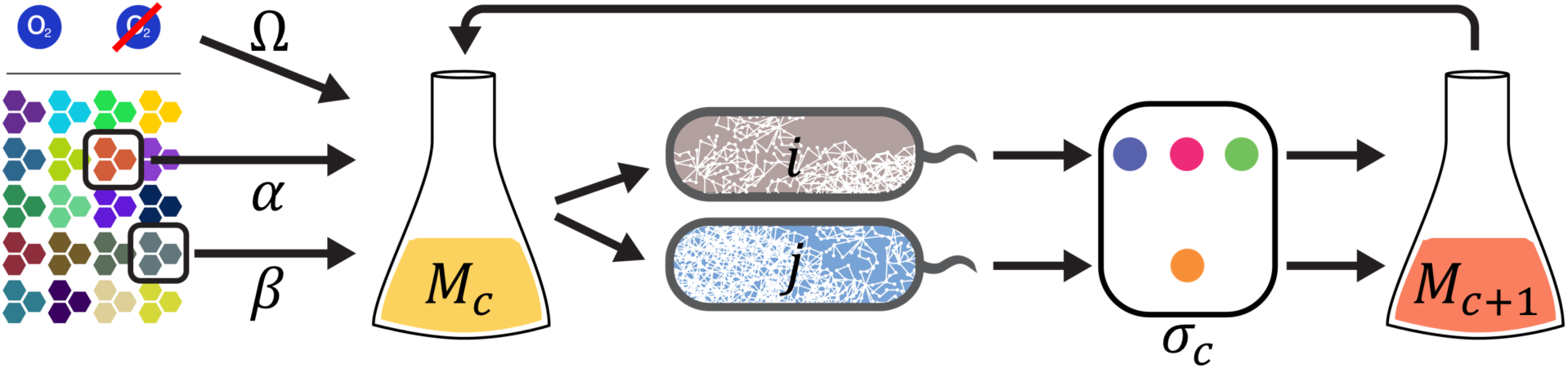
Simplified schematic of example computational pairwise cross-feeding simulation. Simplified schematic of an *in silico* experiment: A growth medium (*M_c_*) containing two carbon sources (*α*, *β*) with or without oxygen (Ω) is provided to genome-scale metabolic models of two microbial organisms (*i*, *j*). If at least one organism grows, any costlessly-secreted metabolites (*σ_c_*) are added to the medium, which is fed back to the organisms. This process is repeated for a series of iterations *c*, and terminates at iteration *c_s_*, defined as the last iteration in which any new metabolites were secreted into the medium.

**Figure 2.**
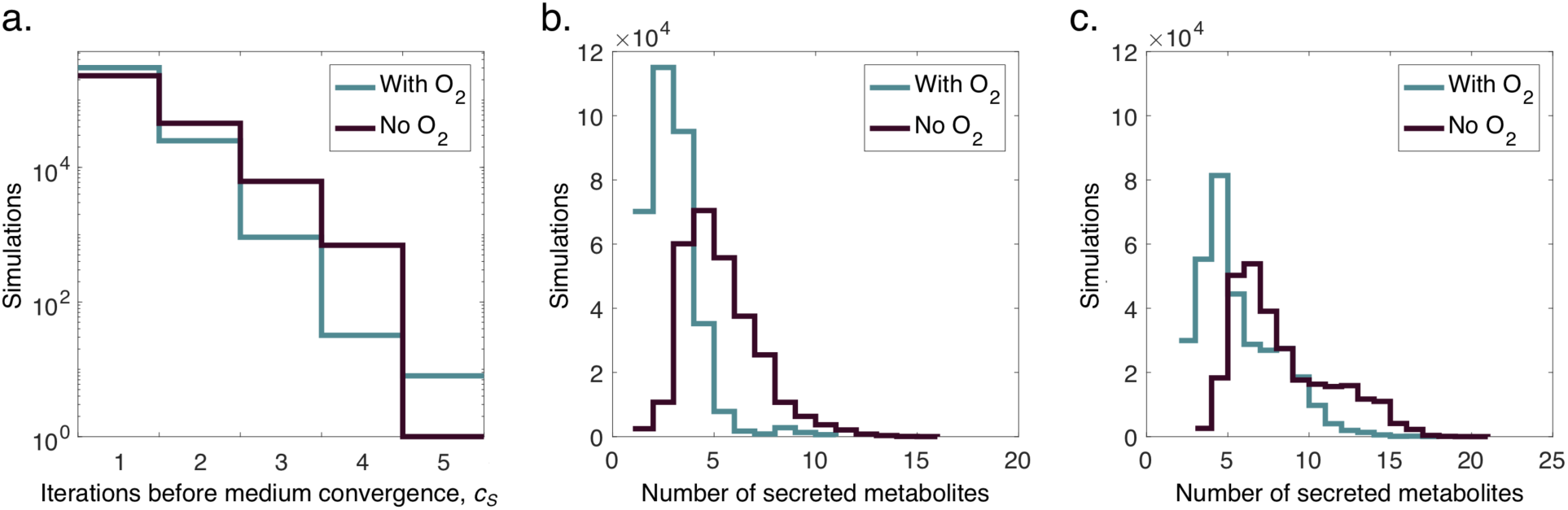
Analysis of costlessly-secreted metabolites in pairwise *in silico* experiments that led to growth of at least one organism. **(a)** Distribution of number of expansions until final medium expansion iteration. **(b)** Distribution of the number of metabolites secreted into the medium by one or both organisms in a pair after one iteration of FBA. **(c)** Distribution of the number of metabolites secreted by one or both organisms after the last iteration of FBA (*c_s_*). The last iteration is defined as the iteration in which no additional metabolites were secreted into the medium. Despite the large variability in number of expansions and number of secreted metabolites, we observe a poor correlation between these distributions, indicating that a simulation resulting in a high number of expansions does not necessarily result in a high number of metabolites being secreted (Figure S3).

In aggregate, our simulations showed a rightward shift in the diversity of metabolites secreted under anoxic conditions when compared to the number secreted when oxygen was present. To understand this effect, we looked at the distribution of the number of metabolites secreted after the first iteration, which is equivalent to growing each organism on its own in the provided medium. This distribution for *c* = 1 was unimodal for both conditions, centered between two and three metabolites with oxygen and around five metabolites without oxygen (Figure 2b). After this first iteration, the maximum number of secreted metabolites was 11 with oxygen and 16 without oxygen. In the anoxic simulations, the central carbon metabolites most commonly secreted after the first iteration were fermentation byproducts such as acetate, formate, succinate, and ethanol. These metabolites were secreted in 87.5%, 74.5%, 25.7%, and 20.2% of growth-yielding simulations respectively. With oxygen, the most commonly secreted central carbon metabolites after the first iteration were formate and acetate, secreted in 46.8% and 18.3% of growth-yielding simulations respectively. We may therefore chiefly attribute the shift between the oxic and anoxic secretion curves to the anoxic export of incompletely-reduced core metabolism intermediates.

In addition to a positive shift observed between anoxic and oxic conditions, our results also show a shift in the quantity of metabolites secreted between the first and last iteration of each computational experiment (Figure 2c). This effect reflects organisms taking up metabolites secreted by themselves or their partner, and secreting different metabolites as a response. After the last medium expansion iteration for all simulations, the total number of secreted metabolites followed similar distributions with a maximum at 18 and 21 metabolites for oxic and anoxic conditions, respectively. This positive shift suggests a response from one or both organisms to a medium iteratively enriched by costless byproducts, which hints at their potential metabolic utility. Principal component analysis (PCA) shows that neither the environment nor the species alone can explain the variability in secretion profiles (Figure S4), suggesting that a combination of both variables accounts for the range in costlessly-secreted products.

### Useful costlessly-secreted byproducts are abundant

Our analysis reveals a broad distribution of metabolically useful compounds secreted without cost in a variety of environmental conditions by most organisms (Figure 3, Figure S5a). Though inorganic compounds such as water and carbon dioxide were, as expected, the most commonly secreted compounds across all simulations, nitrogen-containing compounds such as nitrite, ammonium, urea, and trimethylglycine were secreted in 73.5% of the analyzed cases, suggesting maintenance of an appropriate carbon-to-nitrogen ratio in the cell. We note specifically that nitrite is secreted in fewer than 100 simulations with oxygen, but almost universally in anoxic simulations - a phenomenon previously observed in anaerobic enteric bacteria ^26^. Organic acids make up the second most abundant category of costlessly-secreted byproducts, constituting 23% and 36% of unique metabolites with and without oxygen respectively. Notably, we also observe secretion of nucleotides, peptides, and carbohydrates in a combined 9% and 13% of simulations with and without oxygen respectively. Altogether, this space of secreted metabolites points to a large variety of molecules that can be freely produced, suggesting that costless metabolic secretion may provide substantial degrees of environmental enrichment. This effect becomes magnified considering the relative scarcity of resources provided in our minimal medium, which suggests that costless secretions play a fundamental role in promoting metabolic diversity in natural environments.

**Figure 3.**
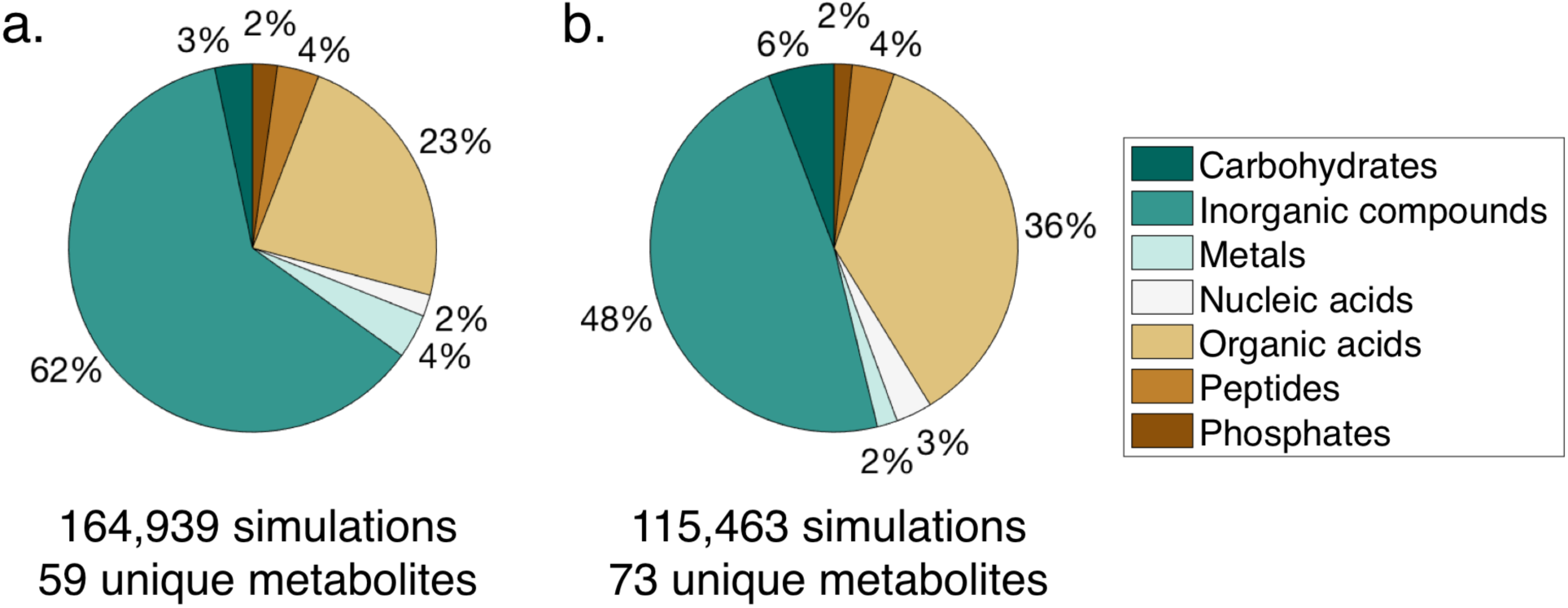
Categorization of metabolites secreted costlessly in oxic (a) and anoxic (b) conditions.

Given the abundance and complexity of secretions from different organisms, as well as the possible ecological connections they may promote, we asked whether specific metabolite secretions were highly correlated. As patterns in environmental modification through secretion have an impact on the species composition of a microbial community ^27^, it becomes important to understand which metabolites co-occur within our set of simulations. To address this question, we performed a Spearman correlation analysis to determine common secretion patterns (Figure S6). In the presence of oxygen, we observe a strong co-occurrence of glycerol, lactate, succinate, malate, and acetate, which correlates with the high frequency of secretion of these carbon-containing compounds (Figure S5a). We also observe positive, but weaker correlations between these metabolites and other central carbon compounds such as fumarate, citrate, and 2-oxoglutarate. Our analysis also points to the simultaneous release of multiple nitrogen-containing compounds, chiefly urea, ammonium, and nitrate. Without oxygen, we observe stronger correlations between secretion of nitrogen-containing compounds and fermentation byproducts. Amino acids also co-occur with high frequency without oxygen, particularly cysteine, methionine, and alanine – which itself is associated with export of proline and glutamine. These patterns are consistent with specific examples of previously studied exometabolomic profiles, including those showing co-secretion of central carbon intermediates in *E. coli* and of amino acids in yeast ^28^, as well as time-dependent patterns of metabolites released simultaneously in soil communities ^29^. In summary, by promoting secretion of a larger number of metabolites across a wide space of conditions, these co-secretion profiles may result in enhanced metabolic enrichment of the environments in our simulation set.

We note that while a potentially useful metabolite can be secreted into the environment by one species, it does not necessarily mean that it will be consumed by a second organism. We place particular importance on this distinction, as any interspecies interaction must also take into account the decision to import a novel metabolite found in the environment. To map this distinction, we examine the space of costless metabolites that are exchanged by each organism across all *in silico* experiments (Figure S5b). Here, the most commonly exchanged organic metabolites were central carbon intermediates, secreted mostly in anoxic conditions. These secretion patterns mirror those of anoxic gut bacteria, which divide the task of digesting complex polysaccharides by exchanging intermediate organic acids ^9,30^. Importantly, we observed that amino acids, secreted chiefly by *S. cerevisiae*, but also in a substantial number of simulations by *S. enterica*, *K. pneumoniae*, and *E. coli*, were among the most highly-exchanged costless metabolites. This phenomenon has been previously documented in relation to overflow metabolism in *S. cerevisiae* ^31^ and *E. coli* ^32,33^, as well as in yeast-bacteria symbioses ^34,35^, and account for exchange in over 10^4^ simulations with and without oxygen in our study. This high prevalence of exchange underscores the metabolic utility of these secreted byproducts, particularly when contrasted with patterns of secretion in which the most commonly released metabolites were of low or no metabolic utility to a partner organism (e.g. water).

### Costless metabolite exchange enhances growth capabilities

Having mapped the space of metabolites that can be secreted costlessly across a large variety of contexts, we asked if these secreted byproducts could directly enable the growth of other organisms. We find that with oxygen, 95,519 *in silico* experiments predicted growth of both organisms in the minimal medium, accounting for 18.2% of all 525,798 oxic simulations (Figure 4a). Under anoxic conditions, only 11.9% of simulations resulted in growth of both organisms in the minimal medium alone. After the organism pairs were allowed to exchange costlessly-secreted metabolites, our algorithm predicted that 31.4% and 22.0% of simulations would result in both species growing with and without oxygen, respectively. This stage of growth, at *c* = *c_s_*, is analogous to both species growing in the presence of each other’s secreted metabolites *in vivo*. This enhanced growth potential in coculture represents a 72.7% increase in growth-supporting environments with oxygen and an 82.5% increase in environments without oxygen, suggesting that exchange of costlessly-secreted metabolites can enable growth of additional organisms in resource-poor environments.

**Figure 4.**
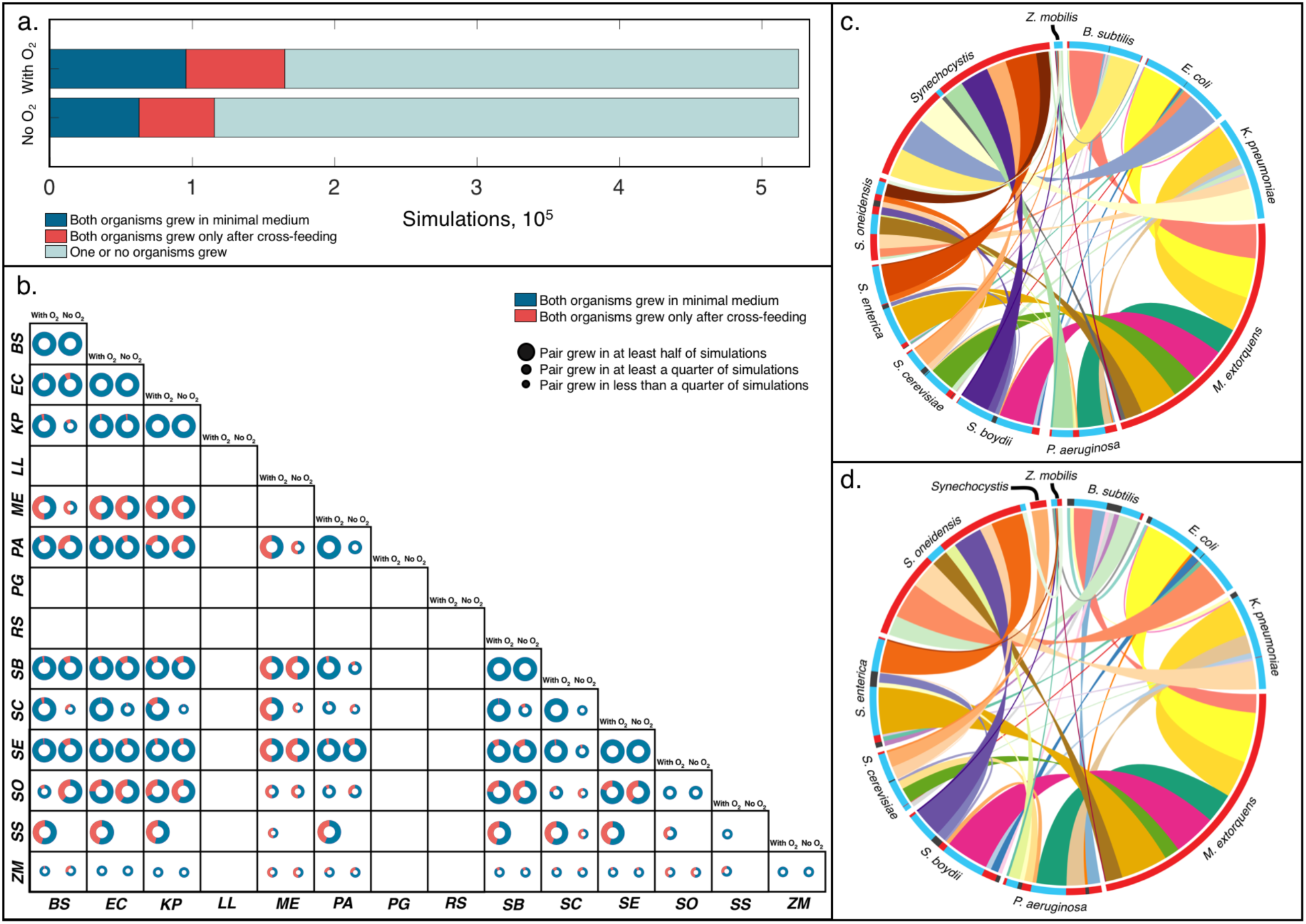
Growth outcomes of pairwise cross-feeding simulations based on organisms and carbon sources. **(a)** Growth outcomes of all *in silico* experiments with and without oxygen, grouped by pairwise growth phenotype. **(b)** Organism-specific growth outcomes. Size of circles represent the relative number of environments in which an organism was able to grow out of 5,774 *in silico* experiments with each partner. Organisms are abbreviated as follows: BS: *B. subtilis;* EC: *E. coli;* KP: *K. pneumoniae; LL: L. lactis;* ME: *M. extorquens;* PA: *P. aeruginosa;* PG: *P. gingivalis;* RS: *R. sphaeroides;* SB: *S. boydii;* SC: *S. cerevisiae;* SE: *S. enterica;* SO: *S. oneidensis;* SS: *Synechocystis;* ZM: *Z. mobilis*. **(c, d)** Frequency of obligate pairwise growth by species in single carbon source simulations for oxic (N = 69,420, c) and anoxic (N = 52,897, d) conditions. Each color ribbon is unique to an individual species pair. Width of ribbons is proportional to the number of experiments in which obligate syntrophy was predicted for each species pair. Radial axis colors represent directionality of exchange: Blue: Organism provided essential metabolites to partner organism in over 75% of simulations; Red: Organism received essential metabolites in over 75% of simulations; Gray: Both organisms gave and received essential nutrients in most simulations.

Though application of our algorithm resulted in a global increase in growth capabilities due to costless metabolite secretion, species-specific growth patterns varied widely across our dataset (Figure 4b). We look specifically at *L. lactis* and *P. gingivalis*, host-associated microbes present in the human gut and oral microbiomes respectively. Both organisms are auxotrophic for a wide range of amino acids and other central metabolites, necessitating dependence on a rich set of metabolic products produced by the host or other commensal microbes. In our simulations, however, these organisms failed to grow in all environments and with all species pairs even after any costless metabolites were secreted. This failure to sustain growth of highly dependent organisms suggests that there is an upper limit to the degree to which costless metabolite production can enable species growth, especially in the minimal environments that were tested. Aside from these extreme cases, our analysis sheds light on the performance of generalist organisms, such as *E. coli*, *K. pneumoniae*, *S. cerevisiae*, and *S. enterica*. These organisms grew in at least half of all tested environmental conditions, in contrast with organisms such as *M. extorquens* or *Z. mobilis*, which exhibited much more limited pairwise growth capabilities. These patterns suggest a greater dependence of these organisms on the metabolic byproducts of their partners, particularly in anoxic conditions. These patterns underscore the importance of not only the number of metabolites secreted, but also of the specific metabolic needs of the receiving organism in determining the contribution of costless metabolites to the growth of a partner.

### Exchange mediated by costless metabolites yields species-specific obligate partnerships

After analyzing general growth outcomes across our entire simulation set, we sought to determine which specific organisms could not grow in our environments without the costless secretions of a partner. This question is of particular interest as co-occurrence in natural communities is widespread ^36–38^ and suggests patterns of species codependence ^39^, potentially providing a mechanistic view into the assembly of complex microbial ecosystems. Our simulations identified a diverse space of codependent organisms, with most species exhibiting at least one case of obligate syntrophy with all others (Figure 4 c, d). Many organisms had balanced distributions of codependence (organism A enabled the growth of organism B in some cases, and organism B enabled the growth of A in others), but the majority of co-dependent relationships were unidirectional. One striking example of this phenomenon is that of cyanobacteria and heterotrophic organisms, with *Synechocystis* (grown here in the absence of light) indicating high degrees of dependence on other organisms. With oxygen, *Synechocystis* was dependent on 9 different organisms across the vast majority of simulations in which it grew with a partner. As all organisms were grown heterotrophically, carbon dioxide and ammonium were the main byproducts that enabled growth of *Synechocystis* in these simulations. Previous studies have confirmed ammonium as the preferred nitrogen source of cyanobacteria ^40–42^, indicating that the ability to fix carbon and consume nitrogen are accurately reflected in the *in silico* metabolic requirements of *Synechocystis*. We also observed that *E. coli*, *B. subtilis*, and *S. cerevisiae*, three species commonly used as model microbial organisms, were more frequently the giving organisms in cases of obligate syntrophy. These pairings not only shed light on the mechanisms behind interspecies codependencies, but may also serve as a map for assembling co-dependent synthetic communities stabilized by costless metabolic exchange.

### Carbon sources exhibit cooperativity in determining growth potential

In addition to characterizing the global space of *in silico* growth phenotypes, we examined how cooperativity of primary carbon sources could enhance growth capabilities in organism pairs. Drawing from techniques used to quantify epistasic interactions ^43^, we defined the cooperativity index C of two carbon sources *α* and *β* as the difference between the number of simulations that result in growth from both carbon sources and the product of the number of simulations that result from single carbon sources. These counts were normalized by the total number of simulations involving the specific pairing of carbon sources being analyzed (represented here by the combinatorial formula 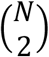), as follows:

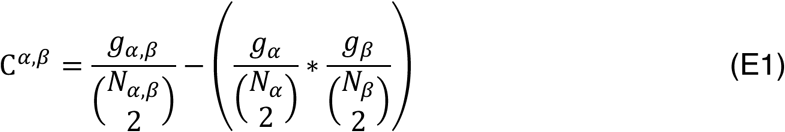

This metric therefore aims to reflect the cooperative potential of each carbon source pair relative to that of each carbon source in isolation. In this way, when averaging a single carbon source over its cooperativity index, we obtain a relative degree to which a carbon source “depends” on another to sustain growth. By framing cooperativity in this context, we observed that simple sugars such as glucose and sucrose had relatively low cooperativity indices, that is, they were able to sustain growth efficiently on their own. In contrast, more complex molecules and dipeptides had higher average cooperativity indices, indicating they performed better in the presence of another carbon source. We grouped these average cooperativity indices through hierarchical clustering (Figure S7) and observed general clustering by carbon source type – especially with sugars and amino acids appearing in distinct groups. This analysis illustrates the nonlinear effects of adding additional nutrients to a minimal medium, underscoring the observed complex metabolite usage patterns in organism pairs.

### Organisms competing for the same carbon source can simultaneously benefit each other through costless secretions

Our analysis so far has examined the contexts in which a metabolite can be secreted costlessly, as well as the potential for these metabolites to promote growth. Based on these insights, we wished to more fundamentally understand these interspecies interactions and how they compare to ecological expectations of cooperation and competition. To do this, we defined six types of possible interactions: non-interaction, commensalism (unidirectional exchange), and mutualism (bidirectional exchange), each with or without competition for a primary carbon source. We chose to decouple competition for nutrients from cooperation via secreted metabolites in order to more fully understand the degree to which the latter can promote organism coexistence despite resource scarcity (Figure 5a). When analyzing our dataset under this framework, we found that competition for one or both carbon sources constituted the majority of the space of all interactions across all simulations (Figure 5b), as previously observed experimentally ^44^. However, these predicted competitive phenotypes were observed to occur simultaneously with potentially beneficial interactions mediated by metabolic byproducts. Here, we found that uni- and bidirectional exchange accounted for a majority of all interactions predicted with and without the presence of oxygen.

**Figure 5.**
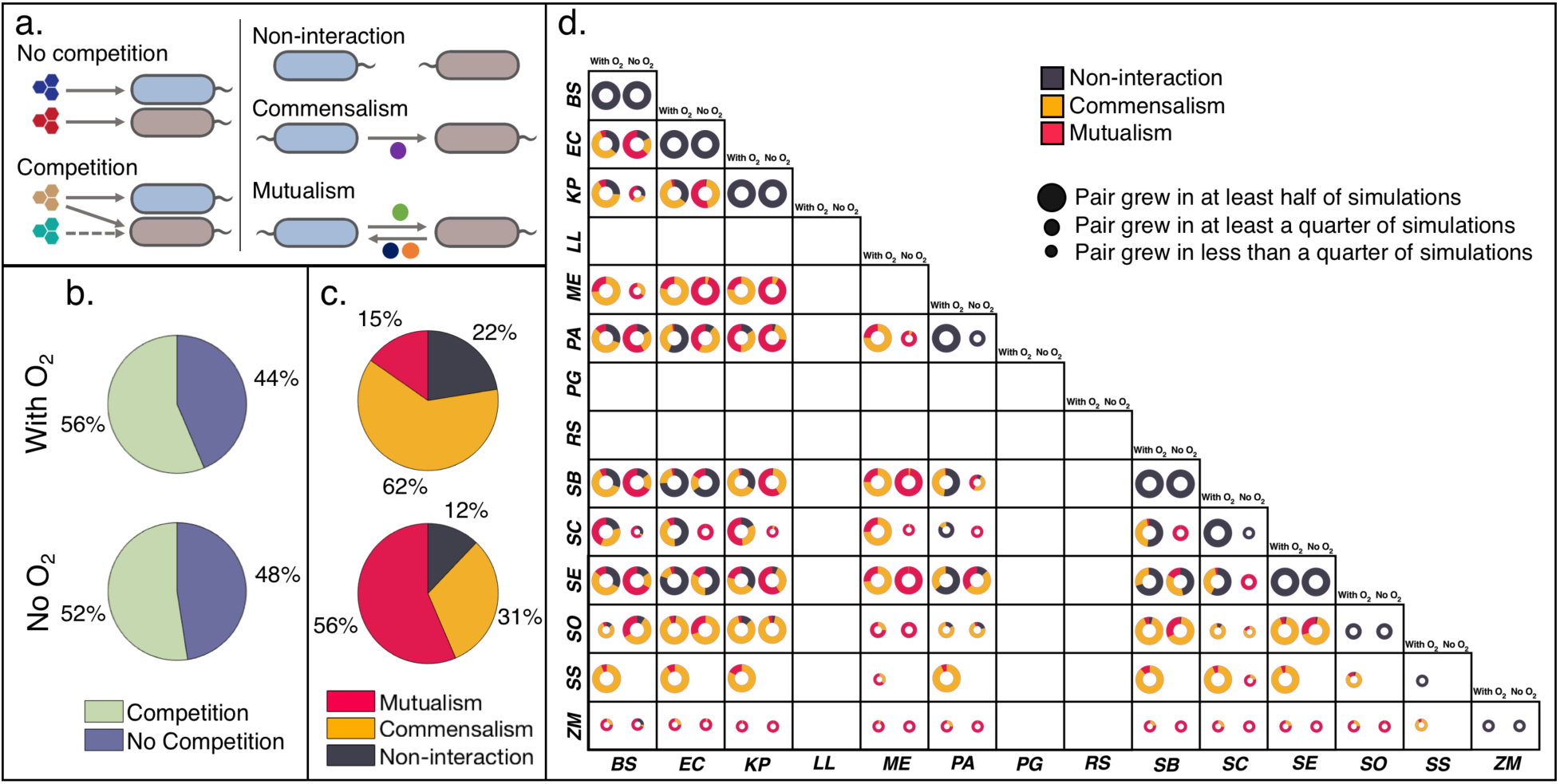
Distribution of metabolic interaction types. **(a)** Schematic representation of interaction types arising from costlessly-secreted metabolites. Competition is defined as both organisms consuming the same carbon source. Commensalism is defined as a unidirectional exchange of one or more costlessly-secreted metabolites, and mutualism is defined as a bidirectional exchange of one or more costlessly-secrete metabolites. **(b)** Overall distributions of competitive/noncompetitive interactions for oxic (out of 164,939 simulations that yielded pairwise growth) and anoxic conditions (out of 115,463 simulations that yielded pairwise growth). **(c)** Overall distributions of general interactions mediated by costless metabolites for oxic and anoxic conditions. These interactions at the level of secreted metabolites exist simultaneously with competition or no competition for a primary carbon source. **(d)** Organism-specific growth outcomes and interaction type distributions. Size of circles represent the relative number of environments in which an organism was able to grow out of 5,774 *in silico* experiments with each partner. Organisms are abbreviated as follows: BS: *B. subtilis;* EC: *E. coli;* KP: *K. pneumoniae; LL: L. lactis;* ME: *M. extorquens;* PA: *P. aeruginosa;* PG: *P. gingivalis;* RS: *R. sphaeroides;* SB: *S. boydii;* SC: *S. cerevisiae;* SE: *S. enterica;* SO: *S. oneidensis;* SS: *Synechocystis;* ZM: *Z. mobilis*.

Our modeling predicted bidirectional interactions to be far more common without oxygen than with oxygen (Figure 5c). We obtained a more fine-grained perspective on costless metabolic interactions by considering the distributions of interaction types by species pairs (Figure 5d). For example, the majority of pairings of *M. extorquens* with *B. subtilis*, *E. coli*, and *K. pneumoniae* exhibited commensal interactions (chiefly with *M. extorquens* receiving). In contrast, the distribution of interactions shifted toward mutualism when oxygen was made unavailable. These patterns were also mirrored in a majority of individual species pairings. As with the positive shift observed in the distributions of secreted metabolites (Figure 2b, c), we may attribute the increased prevalence of mutualistic interactions without oxygen to a greater availability of metabolic byproducts that can contribute to reciprocity. To test this hypothesis, we performed a small subset of “hybrid” *in silico* experiments, where we analyzed the interactions that arose from one species being grown in the presence of oxygen and the other anoxically. We looked at the examples of *E. coli* with *B. subtilis* and *S. enterica*, whose pairwise simulations showed greater amounts of mutualistic interactions without oxygen. When *E. coli* was grown anaerobically but its partner was grown with oxygen, the vast majority of interactions observed were unidirectional, with *E. coli* providing costless metabolites to its partner (Figure 6). When *E. coli* was grown with oxygen, its anoxic partner then provided the majority of metabolites that were exchanged. These intermediate hybrid simulations thus serve as a type of stepping stone, in which an organism grown anoxically can provide a higher number of useful byproducts to its aerobic partner, leading to bidirectional interactions when both are grown without oxygen.

**Figure 6.**
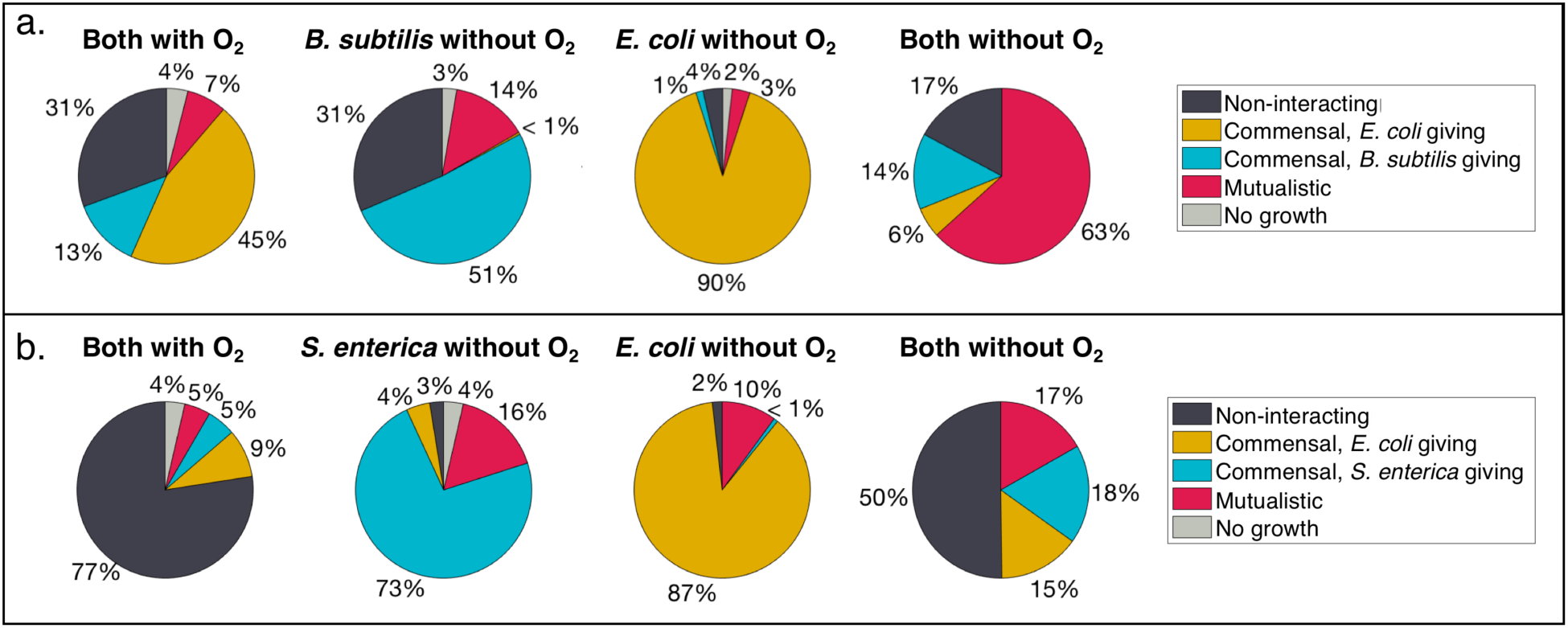
Interaction type distributions from hybrid oxic-anoxic *in silico* experiments for two organism pairs: *E. coli* with *B. subtilis* (a), and *E. coli* with *S. enterica* (b).

### Interaction motifs form a basis for synthetic community assembly

Lastly, we wished to use data generated by our algorithm to understand how multiple simultaneous interactions between two organisms could combine into network patterns (motifs) with different chance of appearance in a community and different dynamical stability properties. In particular, we sought to understand how the competition for common nutrients and the rise of costless exchange could jointly affect the stability of microbial consortia in resource-poor environments. These criteria could also serve as an atlas for guiding the engineering of stable synthetic consortia built off of costless metabolic relationships. As a first step in this analysis, we enumerated possible interaction network motifs based on our three interaction types and competition statuses (Figure 7a). These motifs encompassed all the possible permutations of interactions we identified in our dataset, accounting for non-interaction, commensalism, and mutualism with or without competition. For non-interacting motifs, our simulations predicted an almost exclusive representation of relationships involving competition for a primary carbon source (Figure 7b). The distribution between competitive and non-competitive types motifs was more balanced for commensal and mutualistic interactions, showing a slight preference for interactions involving competition.

**Figure 7.**
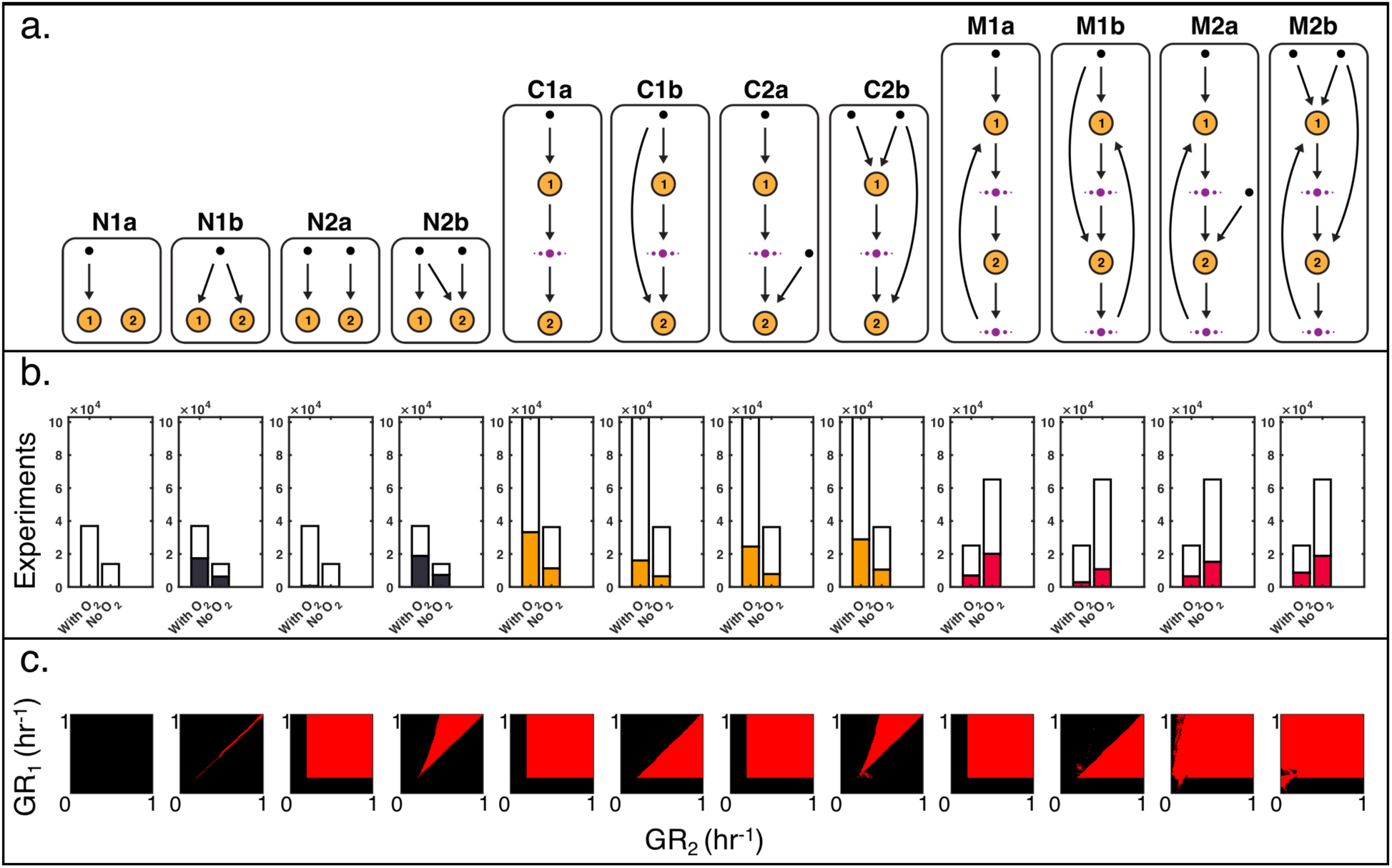
Interaction motif analysis and dynamical modeling of motif stability. **(a)** Schematic representation of specific motif types. Motifs are named according to three features: the interaction type (non-interacting, N; commensal, C; mutualistic, M), the number of carbon sources consumed by the pair (1-2), and competition for a primary carbon source (no competition, a; competition, b). Orange circles denote organisms, black dots denote primary carbon sources, and violet dots indicate any arbitrary number of costlessly-shared metabolites. Arrows indicate direction of metabolite flow. **(b)** Frequency of specific motif types. Height of empty white bars indicate the total number of simulations that exhibited the general motif type (Non-interacting, commensal, mutualistic). Colored bars within indicate the number of the specific motif type (N1a, N1b, etc.). **(c)** Stability space of motifs from dynamical chemostat modeling, as a function of the specific growth rates of the two organisms involved (GR_1_, GR_2_). Red indicates area of stable coculture.

In order to simulate how these interactions could contribute to stable symbioses, we created a dynamical chemostat model of two arbitrary species consuming carbon sources and exchanging costless metabolites according to each motif type (see Methods). By varying the specific growth rates of each species from 0 to 1 hr^−1^, we simulated the growth of the pair under each motif type for 500 hours. If both species were still present at the end of the simulation, we determined the motif type to enable stability at that combination of specific growth rates. We mapped the space of stable species pairs under each motif type, observing that competitive interactions generally have a reduced parameter space for enabling stability (Figure 7c). Notably, though motif N1b was highly prevalent in the costless FBA simulation set, this motif represents classic competitive exclusion and cannot result in long-term stability. In contrast, though complete nutrient-organism orthogonality can yield stability over the whole space of parameters (N2a), this motif was not predicted to occur in the mechanistic simulations. An intermediate case between these two extremes (N2b) is the one in which there is a balance between competition and independence with respect to external carbon source utilization: in this case, which frequently occurs in our dataset, stability is achievable only for a narrow set of parameters.

A marked increase in stability is predicted when costless metabolite exchange is enabled (commensalism and mutualism). For motif C2b, for example, both organisms are competing for a carbon source and organism 1 is providing one or more costless metabolites to organism 2. Our dynamical modeling showed that the growth rate of organism 1 must be greater than that of organism 2 in order for both species to be stable. When feedback was allowed to occur (mutualism), the potential for stability vastly increases across our parameter space. M2a and M2b even allowed for very low specific growth rates for both organisms, indicating a strong dependence on costless metabolites for long-term coexistence.

## DISCUSSION

We have investigated the pairwise growth phenotypes and interactions of 14 diverse microbial species in over 10^6^ computational experiments. We found that resource-poor environments provide the basis for release of a wide variety of useful metabolic products secreted without cost by their producing organism; these costless metabolic products provide, in an oxygen-dependent manner, valuable environmental enrichment, nearly doubling the potential of minimal environments to sustain growth. We further found that exchange of costless metabolites establishes beneficial uni- and bidirectional interspecies interactions, associated with different chance of stability of the ensuing consortia. Overall, both the metabolic capabilities of the organisms and the environmental contexts in which they are grown (particularly oxygen availability) determine which metabolites will be secreted without cost and how these secretions will contribute to interspecies interactions.

Our modeling pipeline represents a novel *in silico* representation of distinct organisms growing in environments progressively enriched by their partner’s secreted byproducts. This iterative medium expansion method provides a useful lens into the emergence of higher-order interactions in microbial communities, allowing us to observe which metabolites are secreted in response to others in a mechanistic fashion. We highlight the utility of applying metabolic modeling to this area, particularly considering the experimental inaccessibility of measuring metabolic secretions, interactions, and stability across all the species and environmental conditions we tested. We observe that, despite allowing organisms to secrete new metabolic products in response to changing medium conditions, the amount and types of metabolites secreted is not enough to sustain prolonged expansion iterations in most cases. This medium expansion distribution hints at an upper limit to higher-order interactions mediated by costless metabolites in microbial ecology.

We nonetheless emphasize that even in the simple, minimal environments we studied, our modeling framework, based on fundamental stoichiometric constraints and metabolic efficiency assumptions, predicts the widespread prevalence of molecular products that are secreted without a metabolic burden and that can benefit other organisms. An important implication of this prediction is that costless metabolites may significantly contribute to enriching environments and sustaining biodiversity, even when organisms are competing for the same primary nutrients. By using costless secretions to cooperate while simultaneously competing for primary nutrients, organisms may escape some of the limitations of pure competition, which has been predicted to limit biodiversity ^45^. This inference could help understanding microbial metabolic dynamics in many different environments, ranging from structured soil communities to large oligotrophic microbial communities, such as those found in the open ocean. This type of exchange, similar to metabolic leakage behind the Black Queen Hypothesis ^17,20^, may contribute to the maintenance of small genomes in resource-poor environments, as the metabolic needs of some organisms can be fulfilled by others. We look specifically at the obligate partnerships predicted by our analysis, which mirror previously-studied codependencies ^9,40,41^. While our algorithm explored only pairs of organisms in coculture, one may wonder whether more complex communities would display qualitatively different features. Our analysis indeed suggests that higher order communities can support growth of highly auxotrophic organisms such as *L. lactis*, *P. gingivalis*, and *R. sphaeroides*: in our pairwise combinations, these organisms did not obtain enough byproducts from any single partner; however, most of the metabolites that these organisms require to grow on a minimal medium were producible separately by multiple species.

Our interaction analysis also provides deeper mechanistic insight into the increased prevalence of mutualistic interactions without oxygen, a phenomenon that has been previously predicted computationally ^46^ and that provides a window into metabolic relationships in environments harboring steep oxygen gradients, such as the human gut ^47^. By carrying out a set of hybrid oxic-anoxic *in silico* experiments, we observed that the additional metabolites secreted anoxically by a facultative anaerobe (e.g. fermentation byproducts) could provide extensive food supply for aerobically growing organisms. This phenomenon has been suggested to play an important role in maintaining equilibrium in communities at oxic-anoxic interfaces in the mammalian gut ^48,49^ and could be the subject of further mechanistic studies.

Although our modeling method considers a wide space of mechanistic constraints in predicting costless metabolic exchange, we acknowledge that secretion patterns and exchange potential are also defined by a variety of other biological factors that fall outside the scope of constraint-based modeling ^50^, such as signaling-based decisions, regulatory states, and thermodynamic gradients induced by metabolite concentrations. Thus, our analysis, in addition to demonstrating the plausibility of widespread costless cross-feeding, could serve as the basis for prioritization of future specific experiments, for which model predictions could be thought of as a null hypothesis against which to compare empirical measurements. Moreover, though our analysis may accurately predict some instances of metabolism-driven synergistic interactions, there may exist experimental barriers (e.g. temperature or pH incompatibilities) to co-culturing some of the organisms in our list, which are not captured in our modeling method. Nonetheless, our mechanistic modeling framework may be applied to finding candidate species-environment pairs that yield mutualistic relationships. Dynamical modeling coupled with these metabolic analyses could then be used to obtain the parameter space most likely to yield desired stable partnerships *in vivo*. Because this approach relies on screening environments that can yield synergy as opposed to engineering individual strains, this approach has the potential to simplify the process of assembling novel synthetic communities ^51^. Our analysis is also easily scalable to a large number of organisms and environments, and could help produce a global atlas of expected, environment-dependent costless secretions and their potential roles in mediating ecological interactions, with applications in understanding and engineering microbiomes.

## METHODS

### Selection and modification of genome-scale metabolic models

A genome-scale metabolic reconstruction was obtained for each of the 14 facultative anaerobic organisms used in the analysis ^52–65^. Genome-scale metabolic models are mathematical representations of an organism’s known metabolic network, which are used to generate mechanistic predictions of growth and resource allocation in a variety of environmental conditions. The process of generating a genome-scale metabolic model has been outlined conceptually ^66–69^ and described procedurally ^70^ by various groups, and generally comprises an automatic generation of a model based on pathway and genome data followed by manual curation by integrating phenotyping, metabolomic, or transcriptomic data ^71^. We note that although an automatically-generated draft metabolic model can be constructed for virtually any organism for which a genome annotation exists, the space of high-quality, experimentally-verified metabolic models that have undergone the manual curation process summarized above is comparatively very small ^72^. This is due to the time and resources needed to complete the curation process, which can span from six months ^70^ to more than ten years for the iteratively-refined model of *E. coli* K-12 ^64^. We nonetheless consider this process to be essential in producing models that can generate the mechanistic cross-feeding predictions detailed here, which rely on verified metabolic capabilities in monoculture.

The models used in this analysis span four taxonomic kingdoms, including representatives from eight bacterial taxa, as well as a variety of primary metabolic strategies (Supplementary Information 1). In addition, these models describe several organisms that are commonly used for *in vivo* studies (*E. coli* K-12, *S. enterica* LT2, etc.), making the resulting costless cross-feeding predictions particularly useful for synthetic ecology experiments and microbial community assembly.

Each model was imported into MATLAB (The MathWorks, Inc., Natick, Massachusetts) using the COnstraint-Based Reconstruction and Analysis (COBRA) Toolbox ^73^, a software platform for constraint-based modeling of metabolic networks. In order to enable *in silico* cross-feeding to be correctly classified, the namespace of all of the metabolic compounds in each of the models was standardized to be internally consistent. This was performed via a computational pipeline with additional manual curation for irregularly-annotated metabolites.

### Computational methodology description and inputs

Our computational method comprises a set of programs written in MATLAB that use Flux Balance Analysis (FBA) to mechanistically define the growth status and metabolic exchange of microbes through costlessly-secreted byproducts. Briefly, FBA is a mathematical method that determines an optimal distribution of metabolic flux through a biochemical network that will maximize a given objective, usually biomass ^74^. An FBA problem is framed in the context of several constraints, namely: (i) *S*, the stoichiometric matrix of dimensions *m* × *n* where *m* is the number of metabolites and *n* is the number of reactions in the model; (ii) *v*, the vector of all reaction fluxes; and (iii) *v_min_* and *v_max_*, flux constraints placed on *v*, defined by enzymatic capacity and experimentally measured uptake rates.

We employ FBA to determine if an organism is able to grow on the *in silico* growth media conditions we define, in addition to which metabolites are taken up and costlessly secreted. We first apply FBA by maximizing for growth and obtaining an optimal growth rate for an organism, 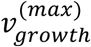. To determine which metabolites are secreted costlessly, we set this growth rate as a minimum for the biomass flux and apply FBA again, recording any metabolites that were secreted. We also apply the additional constraint of minimizing all reaction fluxes across the network to more closely simulate efficient use of the proteome and minimize cycling of metabolites through the network ^75^. Our linear program therefore becomes:

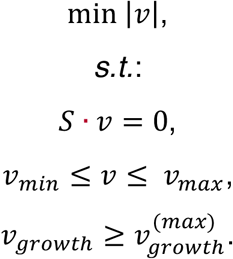

This optimization aims to encompass any enzymatic cost incurred by the organism in synthesizing and exporting any metabolite we deem to be ‘costless.’ During each step in which growth or metabolite absorption and secretion are computed, FBA optimizations are performed separately for each *in silico* organism *i* and *j*, with biomass production set as the objective function while minimizing the sum of the absolute value of *v*. Because we focus on the emergence of potential metabolic exchange through the availability of costlessly-secreted metabolites, our modeling framework purposefully keeps FBA optimizations separate for each model without accounting for spatial or temporal community structure. It is also for this reason that we establish the biomass fluxes of each *in silico* organism as the objective functions to be optimized, as we are concerned with secretion of potentially useful metabolic byproducts that arise out of “selfish” optimal growth. This assumption of maximum growth with proteome optimality is also key for translating these organisms and predictions to *in vivo* synthetic ecologies, where biomass optimization more closely describes the behavior of organisms in batch or continuous culture ^76^.

Our algorithm requires six inputs: 1: a data structure containing the genome-scale metabolic models to be used, 2: a list of carbon sources, 3: the number *N_M_* of *in silico* organisms to be simulated together (for pairwise simulations *N_M_* = 2), 4: the number *N_CS_* of carbon sources to be provided to each simulation, 5: a Boolean variable Ω = {1,0} that specifies if oxygen will be made available to the *in silico* organisms, and 6: a list of metabolites that makes up a simulated base growth medium, *M_min_*. This base medium contains various nitrogen, sulfur, and phosphorus sources, as well as vitamins, ions, and metals needed for growth of the organisms (Supplementary Information 3).

We focused on pairwise species growth with two carbon sources (*N_M_*, *N_CS_* = 2). Although each genome-scale metabolic model we used has been manually curated to reflect *in vivo* metabolic capabilities, very few experiments have been performed to verify FBA-generated predictions for more than a single species ^77,78^. We therefore limit the number of *in silico* species to two, in order to interpret the growth and cross-feeding predictions with greater confidence. This limit also constrains the combinatorial space of the simulations, which grows exponentially and becomes numerically intractable with more models and carbon sources. In addition, limiting simulations to *N_M_* = 2 allows for greater experimental accessibility for assembling synthetic ecologies based on costless metabolite exchange. Our algorithm can nonetheless be applied to any {*N_M_*, *N_CS_* > 0}.

The list of all possible carbon sources was defined primarily from the carbon sources contained in the BIOLOG Phenotyping MicroArray 1 (PM1) plate, which is used for phenotyping and curation of genome-scale metabolic models ^79–81^. The carbon sources we selected are common mono- di- and polysaccharides, all 20 amino acids, dipeptides, and organic acids contained in the PM1 plate. We also supplemented the list with additional carbon sources known to be consumed by the *in silico* organisms, for a total of 108 (Supplementary Information 2).

To permit uptake of the metabolites in the medium, the constraint on the uptake flux bound *v_max_* for each exchange reaction pertaining to a medium metabolite was removed in each of the models *i* and *j*. This bound was fully removed (*v_max_* = 1000 *mmol*/*gDW* ∗ *hr*) for non-limiting medium components, and was set to *v_max_* = 10 *mmol*/*gDW* ∗ *hr* for the growth-limiting carbon sources *α* and *β*. This latter value is drawn from experimentally-estimated uptake rates of sugars by *E. coli* in exponential growth conditions ^64^, and is applied equally to all other species to simulate general availability of the carbon sources in the environment. All other exchange reaction *v_max_* values are set to zero to block uptake of metabolites not in the medium.

### Computing growth, secretion, and cross-feeding

We describe the FBA operations at the core of our algorithm as a function *F* that, given a medium condition *M* and organisms *i* and *j*, outputs the binary growth status *g* of the organisms, as well as the set of metabolites *σ* secreted costlessly by the organisms:

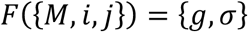

Each *in silico* experiment *E* for a given organism pair with a pair of carbon sources is made up of an initialization step, an expansion step consisting of series of applications of *F*, and a completion step (Figure S2). In the initialization step, two organisms *i* and *j* are selected, and a medium *M*_0_ is defined. *M*_0_ contains the minimal medium *M_min_*, two carbon *α* and *β*, and the variable Ω, which denotes the presence or absence of oxygen. In the expansion step, the function *F* is applied for a series of iterations *c*. In each *F* simulates the growth of both organisms in the current medium condition and returns the Boolean growth statuses *g_c_* = {*g_i_*, *g_j_*} (where *g_i_*, *g_j_* = {0,1}) of both organisms and the set of any costlessly-secreted metabolites, *σ_c_*. To avoid recording metabolites reported to be secreted only as a result of numerical uncertainty in FBA, a minimal lower flux bound of 0.01 *mmol*/*gDW* ∗ *hr* was applied as a cutoff for determining secretion. If at least one organism in the pair grows, the medium is supplemented with *σ_c_*:

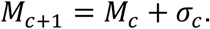

As long as new metabolites continue to be secreted into the medium, that is,

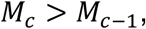

*F* continues to be applied. This stepwise expansion simulates the organisms responding to the costlessly-secreted metabolites being secreted and generating a richer medium. The completion step occurs when no new metabolites are secreted,

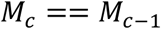

and the final iteration before this stabilization occurs is defined as *c_s_*. Our algorithm therefore carries out individual *in silico* experiments 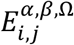, defined as the output resulting from *c_s_* applications of *F* given organisms *i* and *j*, carbon sources *α* and *β*, and the presence or absence Ω of oxygen:

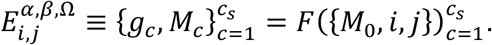

### Dynamical modeling of interaction motifs

We designed a dynamical modeling method to simulate the long-term stability of each pairwise interaction type observed in our *in silico* experiments. We first established a graph theory framework to map each simulation to a specific interaction motif, each of which accounted for the general interaction type (non-interacting, commensal, or mutualistic), the number of carbon sources consumed by the pair, and the competition status for the carbon sources (“a” denotes no competition, “b” denotes competition) (Figure 5a). We next applied a differential equation-based growth model to each specific motif. Since motifs with two carbon sources can be represented by more than one motif topology, we selected one representative topology from these motifs to simplify the space of dynamical modeling simulations. These equations were modeled off Monod dynamics ^82^ and are intended to simulate growth of species in a chemostat, with constant replenishment of medium components. The abundance of each organism *s_i_*, in g/L, is modeled as follows:

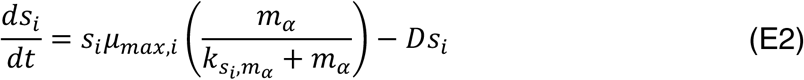

where *μ_max,i_* is the specific growth rate of organism *i* in h^−1^, *m_α_* is the concentration of carbon source *α* in g/L, *k_s_i_, m_α__* is the concentration of *α* at which organism *i* reaches half its maximal growth rate in g/L, and *D* is the chemostat dilution rate in h^−1^. If two carbon sources are present and the organism is determined to take up both by the motif definition, the equation is modified to include a carbon source *β* as follows:

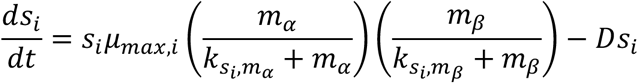

The concentrations of each carbon source are defined as follows:

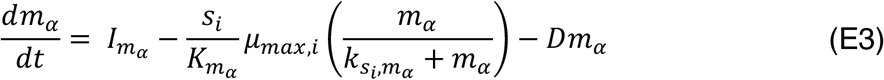

where *I_m_α__* is the nutrient stock concentration for *m_α_* in g/L, and *K_m_α__* is the ratio of nutrient consumed by the organism *i* in g_nutrient_/g_cells_. This equation is modified with an additional term (organism *j*) to simulate competition for *m_α_*.

To simulate metabolic exchange, equations for the abundances of costlessly-produced metabolites (*m̃*) in g/L were defined as follows:

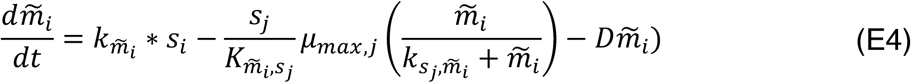

Here, metabolite *m̃*_*i*_ is produced by organism *i* and consumed by organism *j*. *k*_*m̃*_*i*__ is the synthesis rate of the metabolite in hr^−1^, *K*_*m̃*_*i*_, *s_j_*_ is the ratio of metabolite consumed by the population *s_j_* in g_metabolite_/g_cells_, and *k*_*s_j_*, *m̃*_*i*__ is the concentration of metabolite needed for the population *s_j_* to reach half of its maximum growth rate in g/L.

We then combine equations E2-4 to fit the particular motif being modeled (Figure S8). The values of the parameter values are described in Supplementary Information 4 and are based on values reported by Smith ^83^, Balagaddé *et al*. ^84^, and those based on reasonable estimates for resource consumption. For each motif, we vary the specific growth rate of both organisms from 0 to 1 hr^−1^ and run the simulation for 500 hours. If both organism abundances are above 0.05 g/L at the end of the simulation, we determine the motif to be stable at the prescribed growth rates.

## ACKNOWLEDGEMENTS

We thank Dr. Niels Klitgord for pioneering ideas that inspired launch of this work. We are also grateful to David Bernstein, Joshua E. Goldford, Meghan Thommes, Demetrius DiMucci, and all members of the Segrè Lab for helpful discussions. This work was supported by funding from the Defense Advanced Research Projects Agency (Purchase Request No. HR0011515303, Contract No. HR0011-15-C-0091), the U.S. Department of Energy (Grants DE-SC0004962 and DE-SC0012627), the NIH (Grants 5R01DE024468, R01GM121950 and Sub_P30DK036836_P&F), the National Science Foundation (Grants 1457695 and NSFOCE-BSF 1635070), MURI Grant W911NF-12-1-0390, the Human Frontiers Science Program (grant RGP0020/2016), and the Boston University Inter-disciplinary Biomedical Research Office. A.R.P. is supported by a National Academies of Sciences, Engineering, and Medicine Ford Foundation Predoctoral Fellowship and a Howard Hughes Medical Institute Gilliam Fellowship.

## CONTRIBUTIONS

A.R.P. and D.S. designed the research. A.R.P. designed the computational framework, carried out all simulations, and conducted data analysis. M.M. contributed to the generation of standardized genome-scale models. A.R.P. and D.S. wrote the manuscript. All authors read and approved the final manuscript.

## CORRESPONDING AUTHORS

Correspondence to.

## SUPPLEMENTARY FIGURES

**Figure S1.**
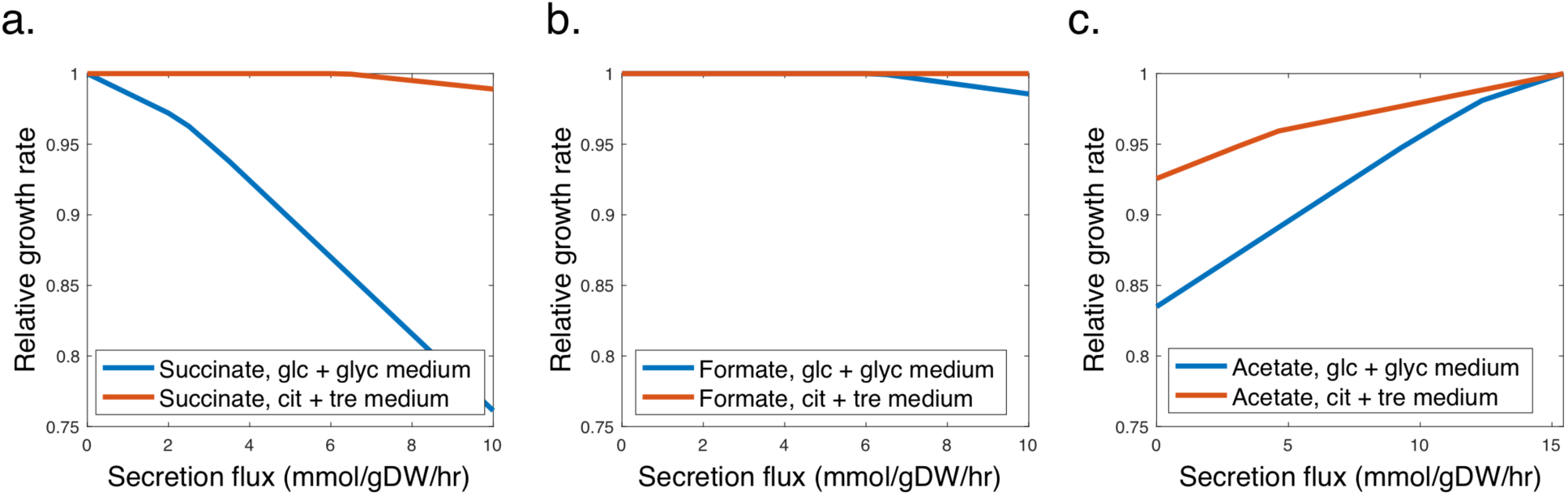
Three modes of *in silico* metabolite secretion by *E. coli* (iJO1366) in anoxic conditions as defined by FBA. What makes a metabolite costless is dependent on the environment. **(a)** Increasing the secretion flux of a ‘costly’ product, such as succinate, imposes a reduction in growth rate when glucose and glycerol are supplied as carbon sources. When the carbon sources are replaced with citrate and trehalose, succinate is secreted without a cost to growth rate. **(b)** With glucose and glycerol as carbon sources, *E. coli* is predicted to have a wide range of fluxes at which formate can be secreted without a cost to its growth rate. Formate would, according to our definition, be secreted ‘costlessly’ by *E. coli* under the applied environmental conditions. **(c)** Some costlessly-secreted metabolites must be secreted at a given rate in order to maximize growth. If an upper bound is placed on acetate secretion, *E. coli* must allocate resources away from biomass in order to cope with its limited ability to secrete fermentation byproducts. Acetate would therefore also be considered a costlessly-secreted metabolite by our definition.

**Figure S2.**
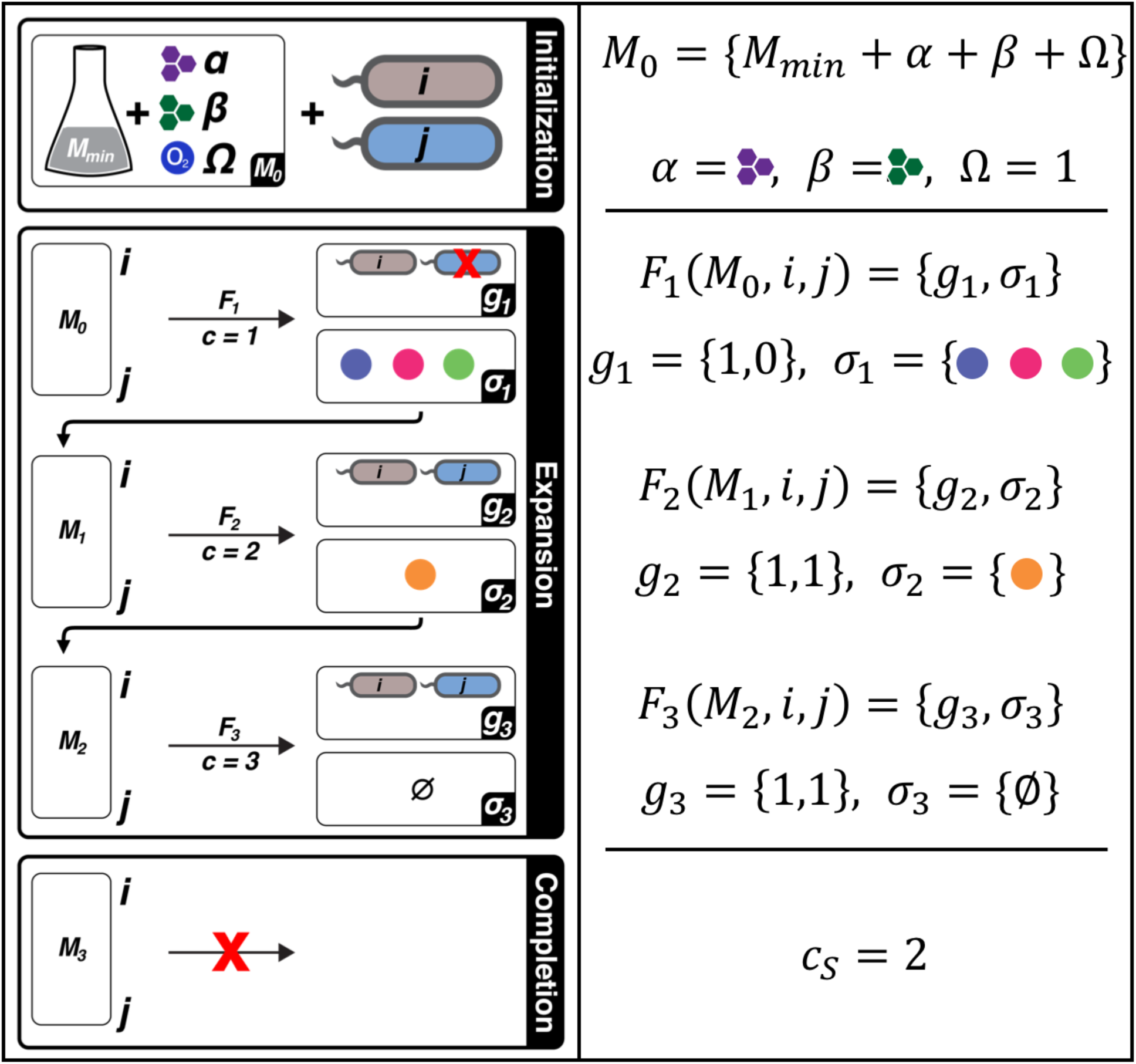
Detailed example of single *in silico* experiment, illustrating three phases. **Initialization:** A minimal medium *M_min_* common to all simulated conditions (composed of salts, metals, vitamins, as well as nitrogen, phosphorous, and sulphur sources) is defined prior to execution of the pipeline. This medium is supplemented with two carbon sources, *α* and *β*.The Boolean variable Ω = {0,1} defines whether or not oxygen is present in the environment. Here, Ω = 1. These together define the initial medium set, *M*_0_. **Expansion:** The function *F* is applied to genome-scale metabolic models of two organisms (*i*, *j*) in a series of iterations, *c*. In each iteration, *F* simulates the growth of both organisms in the current medium condition and returns the Boolean growth statuses *g_c_* = {*g_i_*, *g_j_*} of both organisms and the set of any costlessly-secreted metabolites, *σ_c_*. Here, in the first iteration, *g*_1_ = {1,0} since organism *i* grew but organism *j* did not. Since at least one organism in the pair grew, the medium is updated (*M*_*c*+1_ = *M_c_* + *σ_c_*) and *F* is applied again until no new metabolites are secreted. **Completion:** When no new metabolites are added to the medium, the experiment is complete. The last iteration with any new secreted metabolites is defined as *c_s_*.

**Figure S3.**
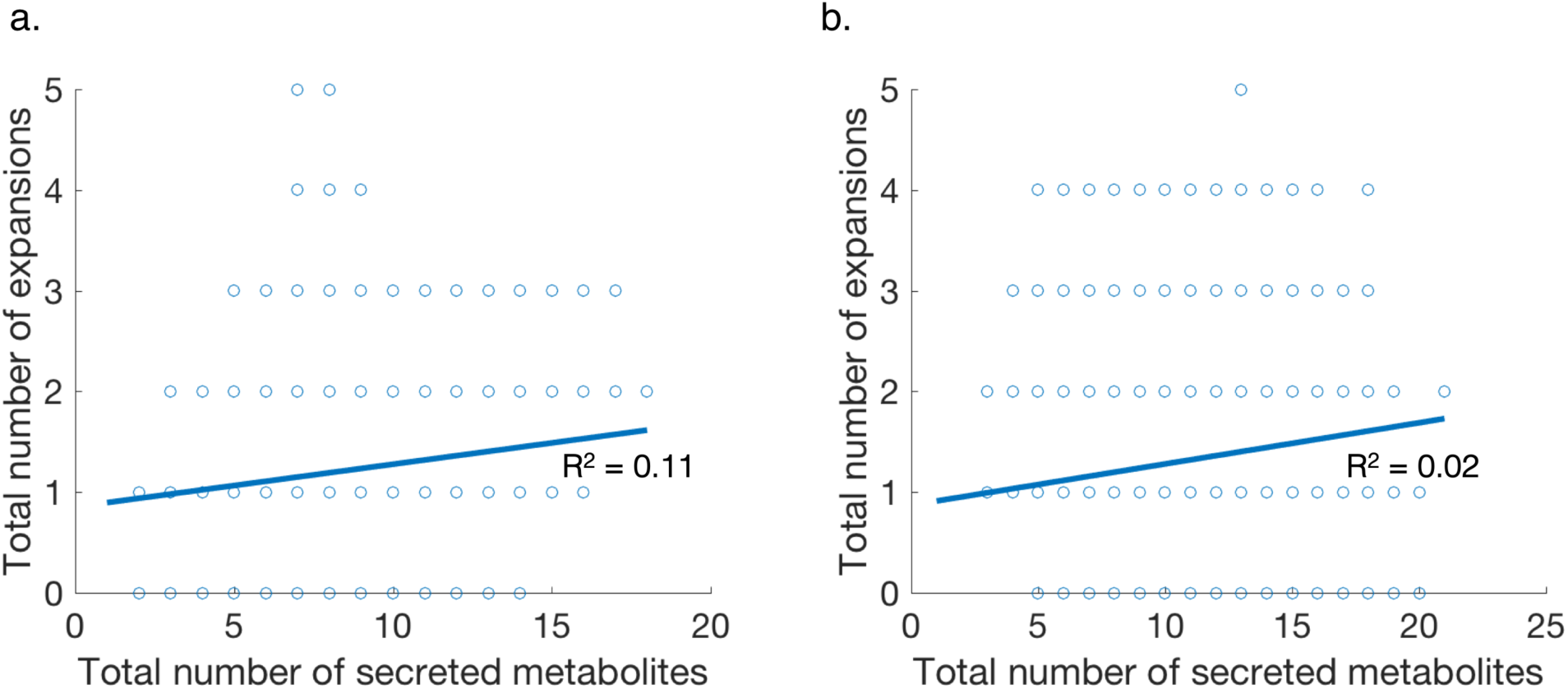
Correlation between total number of metabolites secreted costlessly and the number of expansions in each *in silico* experiment for (a) oxic and (b) anoxic conditions. We observe a poor correlation between number of secreted metabolites and number of expansions in both oxic and anoxic simulations. This lack of correlation suggests a lower rate of metabolite exchange with increasing iterations, with most organisms quickly stabilizing their environment within one or two expansions. With oxygen, for example, only the *K. pneumoniae* and *Synechocystis* pair exhibited more than three medium expansions, with acetate, formate, citrate, and L-malate being the only metabolites secreted at these iterations. These scenarios accounted for only 40 simulations. Without oxygen, there were 697 experiments that reached more than three medium expansions, with 10 organisms being represented. However, this anaerobic set was dominated by the *S. cerevisiae*-*P. aeruginosa* pair, with fermentation byproducts being secreted at late iterations.

**Figure S4.**
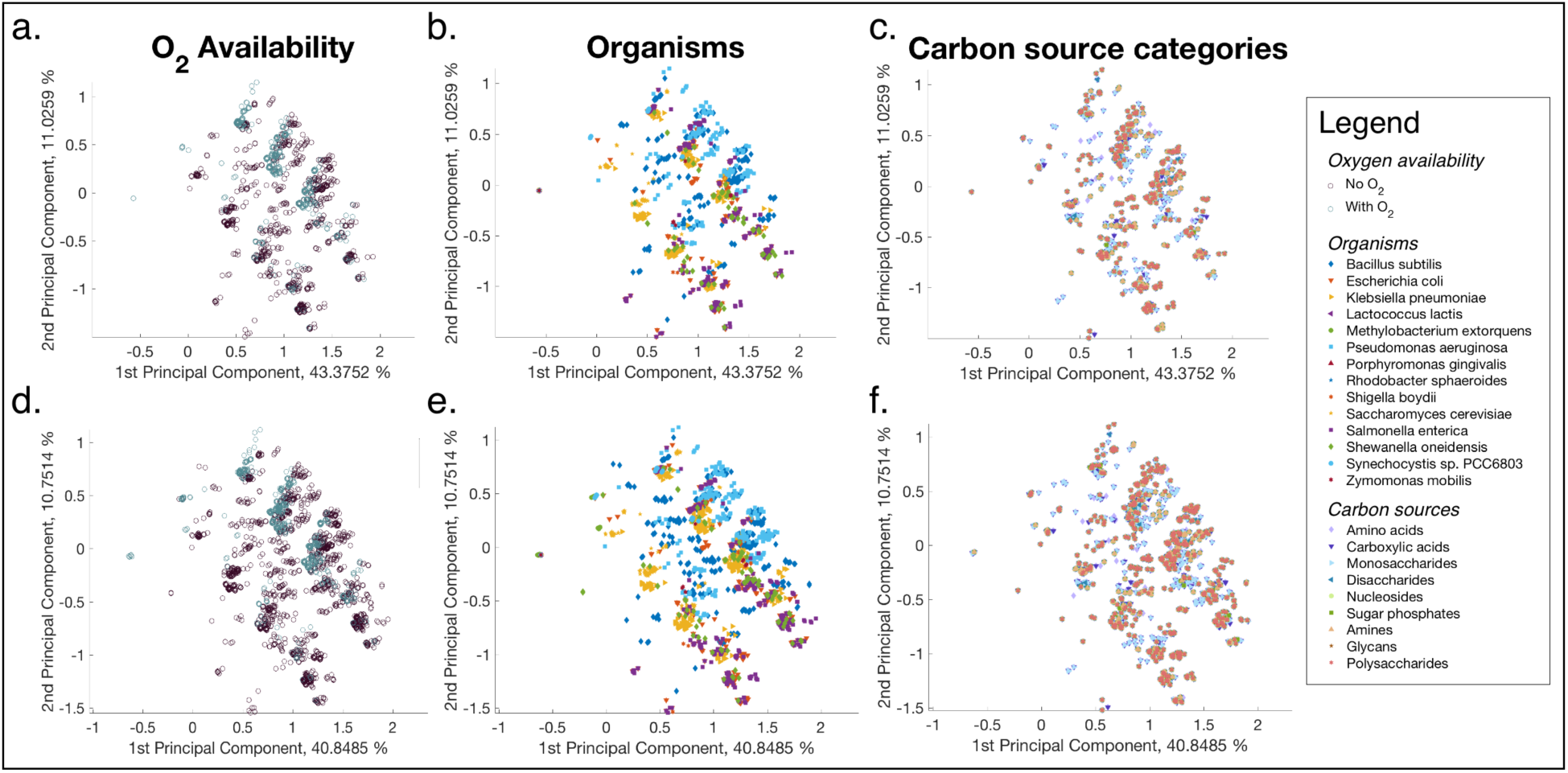
Principal component analysis (PCA) plots for metabolite secretion profiles. To address whether there is one chief contributing factor to patterns of costless metabolite secretion, we carried out principal component analysis (PCA), a dimensionality reduction technique. Each point represents the secreted metabolites of a single organism in one *in silico* experiment. **(a-c)** PCA plots for metabolites secreted before medium expansions (*c =* 1). **(d-f)** PCA plots for secreted metabolites after all medium expansions. Points are clustered by oxygen availability (a, d), organisms (b, e), and carbon source category (c, f).

**Figure S5.**
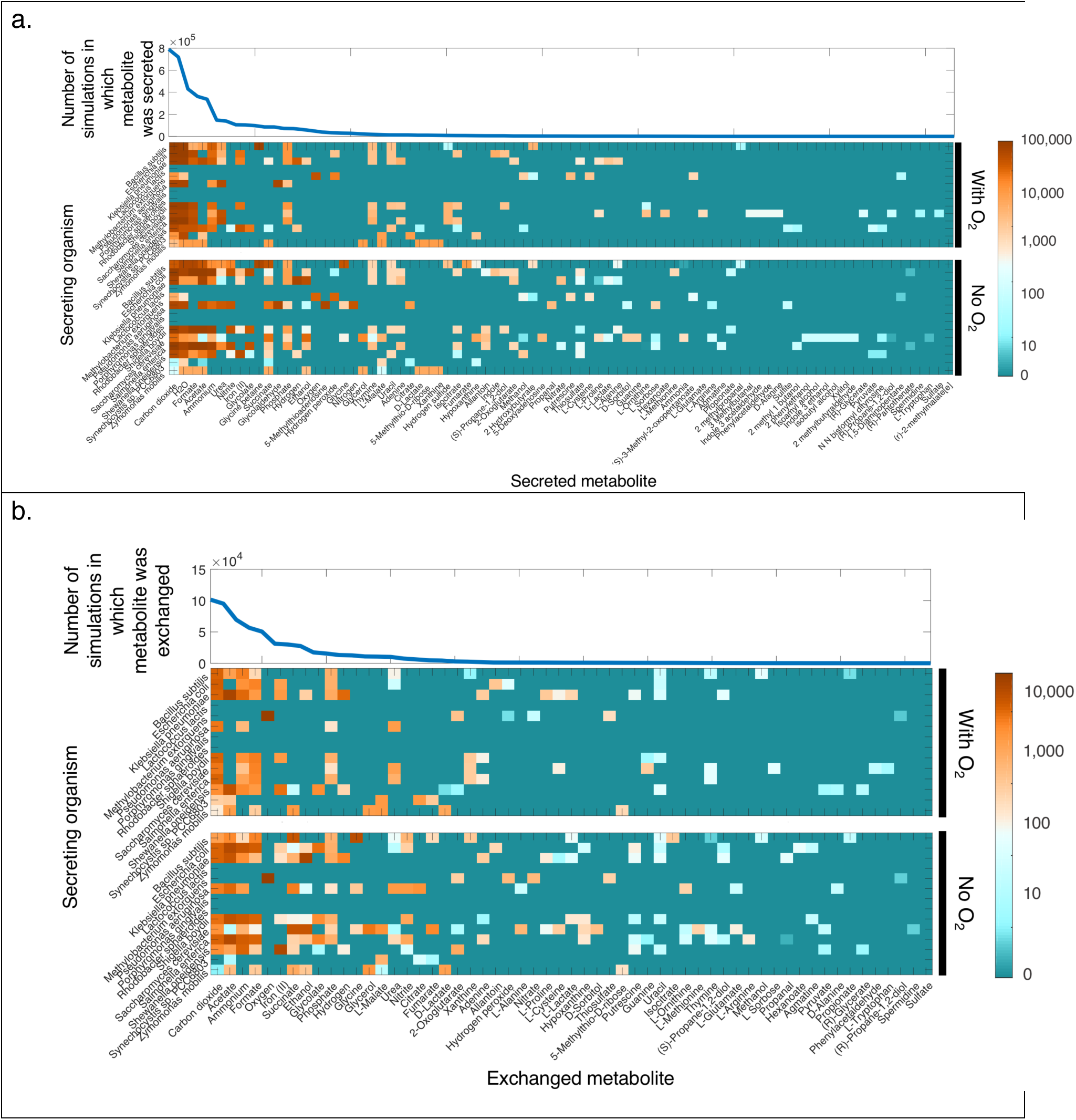
Range of costlessly-secreted and exchanged metabolites. **(a)** Cumulative sum of *in silico* experiments in which metabolite was secreted (top), and sorted heatmap of metabolites secreted in at least one simulation, arranged by secreting organism (bottom). **(b)** Cumulative sum of *in silico* experiments in which each secreted metabolite was taken up by another organism (top), and sorted heatmap of metabolites secreted and taken up in at least one simulation, arranged by secreting organism (bottom).

**Figure S6.**
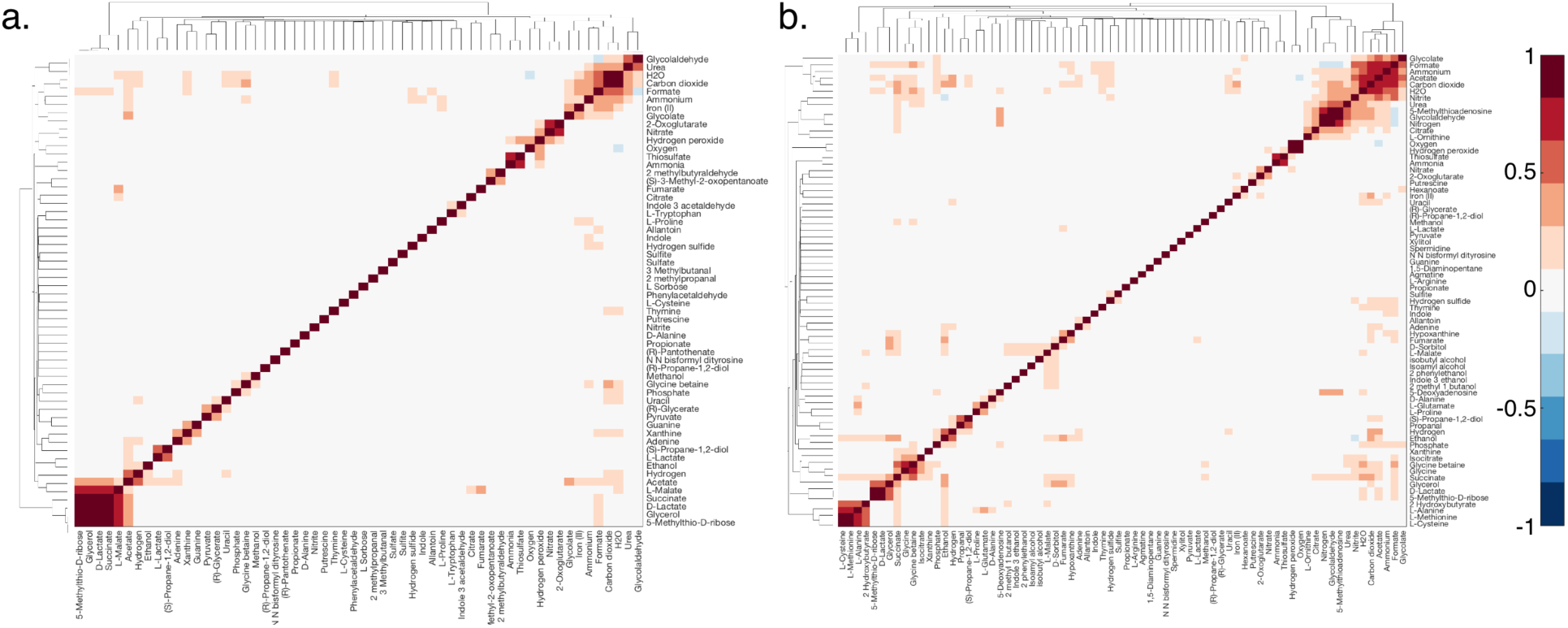
Clustered Spearman correlation of secreted metabolites for (a) oxic and (b) anoxic *in silico* experiments.

**Figure S7.**
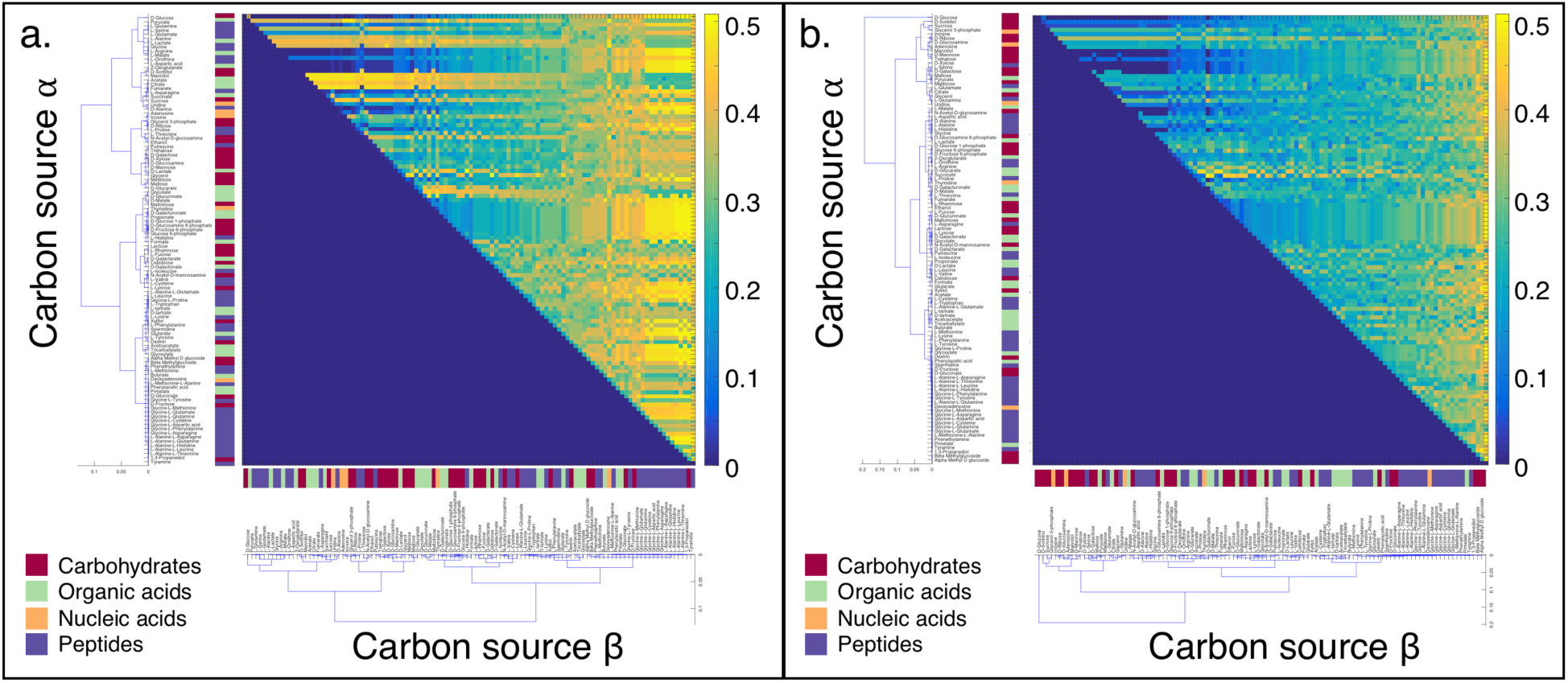
Cooperativity indices of all carbon source pairs in oxic (a) and anoxic (b) conditions, clustered by average carbon source cooperativity index.

**Figure S8.**
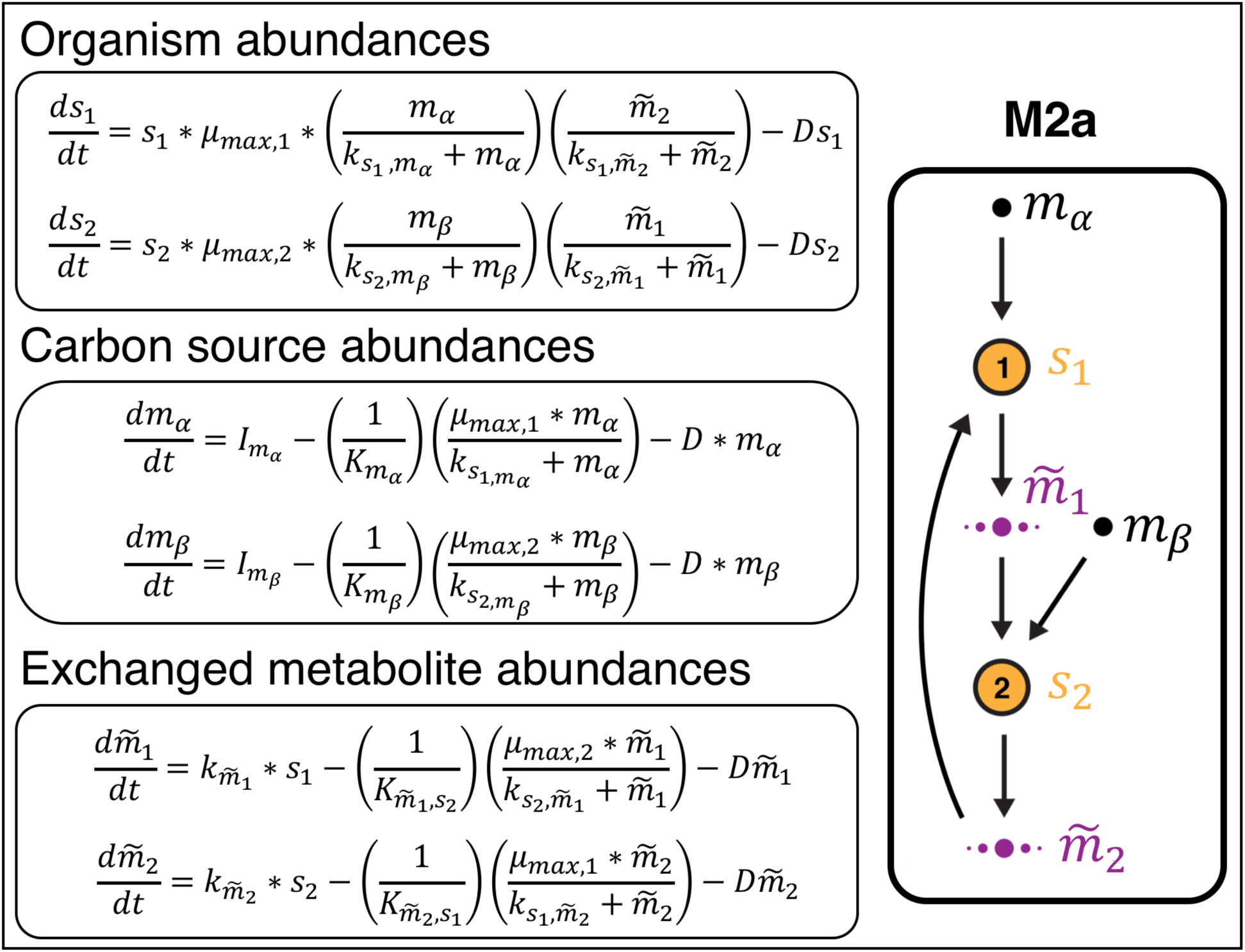
Example of dynamical modeling equations for motif M2a (two carbon sources consumed, no competition, mutualism).

## REFERENCES

1. Welch, D. B. M. & Huse, S. M. Microbial Diversity in the Deep Sea and the Underexplored ‘Rare Biosphere’. Handb. Mol. Microb. Ecol. II Metagenomics Differ. Habitats 243–252 (2011). doi:10.1002/9781118010549.ch24

2. Tecon, R. & Or, D. Biophysical processes supporting the diversity of microbial life in soil. FEMS Microbiol. Rev. 41, 599–623 (2017).

3. Qin, J. et al. A human gut microbial gene catalogue established by metagenomic sequencing. Nature 464, 59–65 (2010).

4. Hardin, G. The competitive exclusion principle. Science 131, 1292–7 (1960).

5. Hutchinson, G. E. The paradox of the plankton. Am. Nat. 91, (1961).

6. Wilson, M. & Lindow, S. E. Coexistence among Epiphytic Bacterial Populations Mediated through Nutritional Resource Partitioning. Appl. Environ. Microbiol. 60, 4468–77 (1994).

7. Inouye, R. S. & Tilman, D. Convergence and Divergence of Old-Field Plant Communities Along Experimental Nitrogen Gradients. Ecology 69, 995–1004 (1988).

8. Kim, H. J., Boedicker, J. Q., Choi, J. W. & Ismagilov, R. F. Defined spatial structure stabilizes a synthetic multispecies bacterial community. Proc. Natl. Acad. Sci. U. S. A. 105, 18188–93 (2008).

9. Morris, B. E. L., Henneberger, R., Huber, H. & Moissl-Eichinger, C. Microbial syntrophy: Interaction for the common good. FEMS Microbiol. Rev. 37, 384–406 (2013).

10. Fildes, P. Production of Tryptophan by Salmonella typhi and Escherichia coli. J. Gen. Microbiol. 15, 636–642 (1956).

11. Goldford, J. E. et al. Emergent Simplicity in Microbial Community Assembly. bioRxiv 205831 (2017). doi:10.1101/205831

12. Ponomarova, O. & Patil, K. R. Metabolic interactions in microbial communities: untangling the Gordian knot. Curr. Opin. Microbiol. 27, 37–44 (2015).

13. Harcombe, W. Novel cooperation experimentally evolved between species. Evolution (N. Y). 64, 2166–72 (2010).

14. Wintermute, E. H. & Silver, P. A. Emergent cooperation in microbial metabolism. Mol. Syst. Biol. 6, 1–7 (2010).

15. Stolyar, S. et al. Metabolic modeling of a mutualistic microbial community. Mol. Syst. Biol. 3, 92 (2007).

16. Vacca, I. Bacterial ecology: Cheaters take advantage. Nat. Rev. Microbiol. 15, 575–575 (2017).

17. Morris, J. J., Lenski, R. E. & Zinser, E. R. The Black Queen Hypothesis : Evolution of Dependencies through Adaptative Gene Loss. MBio 3, 1–7 (2012).

18. Germerodt, S. et al. Pervasive Selection for Cooperative Cross-Feeding in Bacterial Communities. PLoS Comput. Biol. 12, 1–21 (2016).

19. Hoek, M. J. A. va. & Merks, R. M. H. Emergence of microbial diversity due to cross-feeding interactions in a spatial model of gut microbial metabolism. BMC Syst. Biol. 11, 1–18 (2017).

20. Zomorrodi, A. R. & Segrè, D. Genome-driven evolutionary game theory helps understand the rise of metabolic interdependencies in microbial communities. Nat. Commun. 8, 1563 (2017).

21. Sachs, J. L., Mueller, U. G., Wilcox, T. P. & Bull, J. J. The evolution of cooperation. Q. Rev. Biol. 79, 135–60 (2004).

22. West-Eberhard, M. J. The evolution of social behavior by kin selection. Q. Rev. Biol. 50, 1–33 (1975).

23. Connor, R. C. The Benefits of Mutualism: A Conceptual Framework. Biol. Rev. 70, 427–457 (1995).

24. Brown, J. L. Cooperation—A Biologist’s Dilemma. Adv. Study Behav. 13, 1–37 (1983).

25. Orth, J. D., Thiele, I. & Palsson, B. O. What is flux balance analysis? Nat Biotech 28, 245–248 (2010).

26. Tiso, M. & Schechter, A. N. Nitrate reduction to nitrite, nitric oxide and ammonia by gut bacteria under physiological conditions. PLoS One 10, e0119712 (2015).

27. Zelezniak, A. et al. Metabolic dependencies drive species co-occurrence in diverse microbial communities. Proc. Natl. Acad. Sci. 112, 6449–6454 (2015).

28. Paczia, N. et al. Extensive exometabolome analysis reveals extended overflow metabolism in various microorganisms. Microb. Cell Fact. 11, 1–14 (2012).

29. Swenson, T. L., Karaoz, U., Swenson, J. M., Bowen, B. P. & Northen, T. R. Linking soil biology and chemistry in biological soil crust using isolate exometabolomics. Nat. Commun. 9, (2018).

30. Embree, M., Liu, J. K., Al-Bassam, M. M. & Zengler, K. Networks of energetic and metabolic interactions define dynamics in microbial communities. Proc. Natl. Acad. Sci. 112, 15450–15455 (2015).

31. Velasco, I., Tenreiro, S., Calderon, I. L. & André, B. Saccharomyces cerevisiae Aqr1 is an internal-membrane transporter involved in excretion of amino acids. Eukaryot. Cell 3, 1492–503 (2004).

32. Dassler, T., Maier, T., Winterhalter, C. & Bock, A. Identification of a major facilitator protein from Escherichia coli involved in efflux of metabolites of the cysteine pathway. Mol. Microbiol. 36, 1101–1112 (2000).

33. Airich, L. G. et al. Membrane topology analysis of the Escherichia coli aromatic amino acid efflux protein YddG. J. Mol. Microbiol. Biotechnol. 19, 189–97 (2010).

34. Ponomarova, O. et al. Yeast Creates a Niche for Symbiotic Lactic Acid Bacteria through Nitrogen Overflow. Cell Syst. 5, 345–357.e6 (2017).

35. Stadie, J., Gulitz, A., Ehrmann, M. A. & Vogel, R. F. Metabolic activity and symbiotic interactions of lactic acid bacteria and yeasts isolated from water kefir. Food Microbiol. 35, 92–98 (2013).

36. Williams, R. J., Howe, A. & Hofmockel, K. S. Demonstrating microbial co-occurrence pattern analyses within and between ecosystems. Front. Microbiol. 5, 1–10 (2014).

37. HilleRisLambers, J., Adler, P. B., Harpole, W. S., Levine, J. M. & Mayfield, M. M. Rethinking Community Assembly through the Lens of Coexistence Theory. Annu. Rev. Ecol. Evol. Syst. 43, 227–248 (2012).

38. Tilman, D. Resource competition and community structure. (Princeton University Press, 1982).

39. Faust, K. et al. Microbial co-occurrence relationships in the Human Microbiome. PLoS Comput. Biol. 8, (2012).

40. Lindell, D. & Post, A. F. Ecological Aspects of *ntcA* Gene Expression and Its Use as an Indicator of the Nitrogen Status of Marine *Synechococcus* spp. Appl. Environ. Microbiol. 67, 3340–3349 (2001).

41. Glibert, P. M. & Ray, R. T. Different patterns of growth and nitrogen uptake in two clones of marineSynechococcus spp. Mar. Biol. 107, 273–280 (1990).

42. Flores, E. & Herrero, A. in The Molecular Biology of Cyanobacteria 487–517 (Springer Netherlands, 1994). doi:10.1007/978-94-011-0227-8_16

43. Segrè, D., DeLuna, A., Church, G. M. & Kishony, R. Modular epistasis in yeast metabolism. Nat. Genet. 37, 77–83 (2005).

44. Foster, K. R. & Bell, T. Competition, not cooperation, dominates interactions among culturable microbial species. Curr. Biol. 22, 1845–1850 (2012).

45. Ashby, B., Watkins, E., Lourenço, J., Gupta, S. & Foster, K. R. Competing species leave many potential niches unfilled. Nat. Ecol. Evol. 1, 1495–1501 (2017).

46. Heinken, A. & Thiele, I. Anoxic Conditions Promote Species-Specific Mutualism between Gut Microbes In Silico. Appl. Environ. Microbiol. 81, 4049–61 (2015).

47. Espey, M. G. Role of oxygen gradients in shaping redox relationships between the human intestine and its microbiota. Free Radic. Biol. Med. 55, 130–140 (2013).

48. Donaldson, G. P., Lee, S. M. & Mazmanian, S. K. Gut biogeography of the bacterial microbiota. Nat. Rev. Microbiol. 14, 20–32 (2015).

49. He, G. et al. Noninvasive measurement of anatomic structure and intraluminal oxygenation in the gastrointestinal tract of living mice with spatial and spectral EPR imaging. Proc. Natl. Acad. Sci. U. S. A. 96, 4586–91 (1999).

50. Bordbar, A., Monk, J. M., King, Z. A. & Palsson, B. O. Constraint-based models predict metabolic and associated cellular functions. Nat. Rev. Genet. 15, 107–120 (2014).

51. Lindemann, S. R. et al. Engineering microbial consortia for controllable outputs. ISME J. 10, 2077–2084 (2016).

52. Mazumdar, V., Snitkin, E. S., Amar, S. & Segrè, D. Metabolic network model of a human oral pathogen. J. Bacteriol. 91, 74–90 (2009).

53. Motamedian, E., Saeidi, M. & Shojaosadati, S. A. Reconstruction of a charge balanced genome-scale metabolic model to study the energy-uncoupled growth of Zymomonas mobilis ZM1. Mol. BioSyst. 12, 1241–1249 (2016).

54. Peyraud, R. et al. Genome-scale reconstruction and system level investigation of the metabolic network of Methylobacterium extorquens AM1. BMC Syst. Biol. 5, (2011).

55. Imam, S. et al. IRsp1095: A genome-scale reconstruction of the Rhodobacter sphaeroides metabolic network. BMC Syst. Biol. 5, 116 (2011).

56. Flahaut, N. A. L. et al. Genome-scale metabolic model for Lactococcus lactis MG1363 and its application to the analysis of flavor formation. Appl. Microbiol. Biotechnol. 97, 8729–8739 (2013).

57. Pinchuk, G. E. et al. Constraint-Based Model of Shewanella oneidensis MR-1 Metabolism: A Tool for Data Analysis and Hypothesis Generation. PLoS Comput. Biol. 6, e1000822 (2010).

58. Liao, Y. C. et al. An experimentally validated genome-scale metabolic reconstruction of Klebsiella pneumoniae MGH 78578, iYL1228. J. Bacteriol. 193, 1710–1717 (2011).

59. Nogales, J., Gudmundsson, S., Knight, E. M., Palsson, B. O. & Thiele, I. Detailing the optimality of photosynthesis in cyanobacteria through systems biology analysis. Proc. Natl. Acad. Sci. 109, 2678–2683 (2012).

60. Monk, J. M. et al. Genome-scale metabolic reconstructions of multiple Escherichia coli strains highlight strain-specific adaptations to nutritional environments. Proc. Natl. Acad. Sci. 110, 20338–20343 (2013).

61. Zomorrodi, A. R. & Maranas, C. D. Improving the iMM904 S. cerevisiae metabolic model using essentiality and synthetic lethality data. BMC Syst. Biol. 4, 178 (2010).

62. Thiele, I. et al. A community effort towards a knowledge-base and mathematical model of the human pathogen Salmonella Typhimurium LT2. BMC Syst. Biol. 5, 8 (2011).

63. Oberhardt, M. A., Puchałka, J., Fryer, K. E., Martins dos Santos, V. A. P. & Papin, J. A. Genome-scale metabolic network analysis of the opportunistic pathogen Pseudomonas aeruginosa PAO1. J. Bacteriol. 190, 2790–803 (2008).

64. Orth, J. D. et al. A comprehensive genome-scale reconstruction of Escherichia coli metabolism--2011. Mol. Syst. Biol. 7, 535 (2011).

65. Henry, C. S., Zinner, J. F., Cohoon, M. P. & Stevens, R. L. iBsu1103: a new genome-scale metabolic model of Bacillus subtilis based on SEED annotations. Genome Biol. 10, R69 (2009).

66. Reed, J. L., Famili, I., Thiele, I. & Palsson, B. O. Towards multidimensional genome annotation. Nat. Rev. Genet. 7, 130–141 (2006).

67. Feist, A. M., Herrgård, M. J., Thiele, I., Reed, J. L. & Palsson, B. Ø. Reconstruction of biochemical networks in microorganisms. Nat. Rev. Microbiol. 7, 129–143 (2008).

68. Price, N. D., Papin, J. A., Schilling, C. H. & Palsson, B. O. Genome-scale microbial in silico models: the constraints-based approach. Trends Biotechnol. 21, 162–169 (2003).

69. Durot, M., Bourguignon, P.-Y. & Schachter, V. Genome-scale models of bacterial metabolism: reconstruction and applications. FEMS Microbiol. Rev. 33, 164–190 (2009).

70. Thiele, I. & Palsson, B. O. A protocol for generating a high-quality genome-scale metabolic reconstruction. Nat. Protoc. 5, (2010).

71. Oh, Y.-K., Palsson, B. O., Park, S. M., Schilling, C. H. & Mahadevan, R. Genome-scale reconstruction of metabolic network in Bacillus subtilis based on high-throughput phenotyping and gene essentiality data. J. Biol. Chem. 282, 28791–9 (2007).

72. King, Z. A. et al. BiGG Models: A platform for integrating, standardizing and sharing genome-scale models. Nucleic Acids Res. 44, D515–D522 (2016).

73. Becker, S. A. et al. Quantitative prediction of cellular metabolism with constraint-based models: the COBRA Toolbox. Nat Protoc 2, (2007).

74. Orth, J. D., Thiele, I. & Palsson, B. Ø. What is flux balance analysis? Nat. Biotechnol. 28, 245–248 (2010).

75. Holzhütter, H.-G. The principle of flux minimization and its application to estimate stationary fluxes in metabolic networks. Eur. J. Biochem. 271, 2905–2922 (2004).

76. Lewis, N. E. et al. Omic data from evolved E. coli are consistent with computed optimal growth from genome-scale models. Mol. Syst. Biol. 6, 390 (2010).

77. Harcombe, W. R. et al. Metabolic resource allocation in individual microbes determines ecosystem interactions and spatial dynamics. Cell Rep. 7, 1104–1115 (2014).

78. Zomorrodi, A. R. & Segrè, D. Synthetic Ecology of Microbes: Mathematical Models and Applications. J. Mol. Biol. (2015). doi:10.1016/j.jmb.2015.10.019

79. Reed, J. L. et al. Systems approach to refining genome annotation. Proc. Natl. Acad. Sci. 103, 17480–17484 (2006).

80. Oberhardt, M. A., Puchałka, J., Martins dos Santos, V. A. P. & Papin, J. A. Reconciliation of Genome-Scale Metabolic Reconstructions for Comparative Systems Analysis. PLoS Comput. Biol. 7, e1001116 (2011).

81. Henry, C. S. et al. High-throughput generation, optimization and analysis of genome-scale metabolic models. Nat. Biotechnol. 28, 977–982 (2010).

82. Monod, J. The Growth of Bacterial Cultures. Annu. Rev. Microbiol. 3, 371–394 (1949).

83. Smith, H. L. Bacterial growth. (2006).

84. Balagaddé, F. K. et al. A synthetic Escherichia coli predator-prey ecosystem. Mol. Syst. Biol. 4, 187 (2008).

